# WRAD core perturbation impairs DNA replication fidelity promoting immunoediting in pancreatic cancer

**DOI:** 10.1101/2024.10.21.619543

**Authors:** Francesca Citron, I-Lin Ho, Chiara Balestrieri, Zhaoliang Liu, Er-Yen Yen, Luca Cecchetto, Luigi Perelli, Li Zhang, Luis Castillo Montanez, Nicholas Blazanin, Charles A. Dyke, Rutvi Shah, Sergio Attanasio, Sanjana Srinivasan, Ko-Chien Chen, Ziheng Chen, Iolanda Scognamiglio, Nhung Pham, Hania Khan, Shan Jiang, Jing Pan, Ben Vanderkruk, Cecilia S. Leung, Mahinur Mattohti, Kunal Rai, Yanshuo Chu, Linghua Wang, Sisi Gao, Angela K. Deem, Alessandro Carugo, Huamin Wang, Wantong Yao, Giovanni Tonon, Yun Xiong, Philip L. Lorenzi, Chiara Bonini, M. Anna Zal, Brad G. Hoffman, Tim Heffernan, Virginia Giuliani, Collene R. Jeter, Yonathan Lissanu, Giannicola Genovese, Mauro Di Pilato, Andrea Viale, Giulio F. Draetta

**Affiliations:** Department of Genomic Medicine, The University of Texas MD Anderson Cancer Center, Houston, TX 77030, USA; Center for Omics Sciences, IRCCS San Raffaele Scientific Institute, Milan 20132, Italy; Experimental Hematology Unit, IRCCS San Raffaele Scientific Institute, Milan 20132, Italy; MD Anderson UTHealth Graduate School of Biomedical Sciences, Houston, TX 77030, USA; Department of Genitourinary Oncology, The University of Texas MD Anderson Cancer Center, Houston, TX 77030, USA; Department of Immunology, The University of Texas MD Anderson Cancer Center, Houston, TX 77030, USA; Department of Thoracic and Cardiovascular Surgery, The University of Texas MD Anderson Cancer Center, Houston, TX 77030, USA; TRACTION, The University of Texas MD Anderson Cancer Center, Houston, TX 77030, USA; Department of Cancer Biology, The University of Texas MD Anderson Cancer Center, Houston, TX 77030, USA; Department of Surgery, University of British Columbia, Vancouver, BC, V5Z 4E3, Canada; Diabetes Research Group, British Columbia Children’s Hospital Research Institute, 950 West 28th Avenue, Vancouver, BC, V5Z 4H4, Canada; Department of Cell and Developmental Biology, University of Pennsylvania, Perelman School of Medicine, Philadelphia, PA 19194, USA; Immunobiology and Transplant Science Center and Department of Surgery, Houston Methodist Hospital, Houston, TX 77030, USA, Department of Epigenetics and Molecular Carcinogenesis, Division of VP, Research; Department of Oncology, IRBM Spa, Via Pontina Km 30 600, Pomezia, Rome 00071, Italy; Department of Pathology, Division of Pathology/Lab Medicine, The University of Texas MD Anderson Cancer Center, Houston, TX; Department of Translational Molecular Pathology, The University of Texas MD Anderson Cancer Center, Houston, TX 77030, USA; Vita-Salute San Raffaele University, Milan 20132, Italy; Proteomics Core Facility, Department of Bioinformatics and Computational Biology, The University of Texas MD Anderson Cancer Center, Houston, TX 77030, USA; Department of Epigenetics and Molecular Carcinogenesis, Division of VP, Research, The University of Texas MD Anderson Cancer Center, Houston, TX 77030, USA

**Keywords:** Pancreatic cancer, WRAD core, DPY30, DNA replication, chromosomal instability, immunoediting, immunotherapy

## Abstract

It is unclear how cells counteract the potentially harmful effects of uncoordinated DNA replication in the context of oncogenic stress. Here, we identify the WRAD (WDR5/RBBP5/ASH2L/DPY30) core as a modulator of DNA replication in pancreatic ductal adenocarcinoma (PDAC) models. Molecular analyses demonstrated that the WRAD core interacts with the replisome complex, with disruption of DPY30 resulting in DNA re-replication, DNA damage, and chromosomal instability (CIN) without affecting cancer cell proliferation. Consequently, in immunocompetent models, DPY30 loss induced T cell infiltration and immune-mediated clearance of highly proliferating cancer cells with complex karyotypes, thus improving anti-tumor efficacy upon anti-PD-1 treatment. In PDAC patients, DPY30 expression was associated with high tumor grade, worse prognosis, and limited response to immune checkpoint blockade. Together, our findings indicate that the WRAD core sustains genome stability and suggest that low intratumor DPY30 levels may identify PDAC patients who will benefit from immune checkpoint inhibitors.

## Introduction

The occurrence of aneuploidy, a condition characterized by a deviation from the normal chromosome number, is differentially sensed in normal and tumor cells (1). In normal cells, aneuploidy triggers molecular checkpoint activation and immune cells recruitment for their own destruction (2, 3). In contrast, in transformed cells, aneuploidy modulates the expression of cancer driver genes (4), favors tumor-intrinsic heterogeneity, phenotypic plasticity (1), and induces genomic instability (5), which can fuel tumor evolution (6, 7). While cell-intrinsic mechanisms have been implicated in contributing to genomic instability in tumors (8, 9), the molecular mechanisms determining the evolutionary trajectory of genomic instability in cancer cells are poorly understood.

To ensure faithful genome maintenance, precise replication of the entire genome occurs only once per cell cycle through a highly conserved and tightly regulated process that begins with the licensing of DNA replication origins (10). Replication licensing is restricted to the G1 phase of the cell cycle and is defined by the loading of the pre-replicative complex (pre-RC), which comprises the origin of replication complex (ORC1-6 complex), Cdt1 (chromatin licensing and DNA replication factor 1), Cdc6 (cell division cycle 6), and DNA helicase (minichromosome maintenance complex; MCM), onto DNA (11–13). As cells progress through the division cycle, DNA replication origin firing is regulated by the phosphorylation of the pre-RC by CDKs and DKK (Cdc7/Dbf4 kinase), which in turn recruit Cdc45 and GINS as accessory factors to fully initiate DNA replication (10, 13–16).

Inaccurate genome duplication, including under- or over-replication of DNA in late S or G2/M phase, can trigger deleterious consequences, such as DNA damage, chromosomal instability (CIN), and aneuploidy. This ultimately leads to aberrant genome architecture (17–19), dysplasia and malignant transformation when surveillance mechanisms fail to clear the damaged cells (12, 20–22). Although the mechanisms that prevent uncoordinated DNA replication in normal cells are well characterized (10, 13, 23–25), it remains unclear how cancer cells limit unscheduled DNA replication, especially given its increased frequency in the context of aberrant oncogenic signaling (e.g. *KRAS* mutation), loss of cell cycle checkpoints (e.g. *TP53*, *CDKN2A*), and cumulative alterations in epigenetic regulators (e.g. *KMT2B*, *KMT2C*, *KMT2D*, *SETD1A*, and *SETD1B*) – all of which are commonly observed in tumors such as pancreatic ductal adenocarcinoma (PDAC) (26–30).

Previously, we discovered that multiple members of the COMPASS complex (WDR5, ASH2L, Menin) are critical for PDAC tumorigenesis and cancer progression in *KRAS/TP53*-mutated backgrounds (31, 32). In mammals, COMPASS complexes are composed of one of six methyltransferase catalytic subunits of the SET1/MLL family (KMT2A, KMT2B, KMT2C, KMT2D, SET1A, SET1B) and the WRAD core (WDR5, RbBP5, ASH2L, DPY30). The full complement of these proteins is required for efficient histone-3-lysine-4 (H3K4) methylation as well as the regulation of transcriptional initiation and elongation of target genes (33–36).

While WDR5 has been extensively characterized as a core scaffolding component of SET1/MLL methyltransferases and demonstrated to be essential for gene transcription, DPY30 is relatively understudied compared to other WRAD components. It is known that DPY30 must bind ASH2L to enable histone H3K4 trimethylation (37), with loss of DPY30 expression leading to global reductions in H3K4 trimethylation levels in multiple model systems (38–45). Studies in yeast suggest that, within the WRAD core, DPY30 may stabilize the COMPASS complex to ensure the correct positioning of H3K4 methylation at promoters, especially for low-transcribed genes. Indeed, in human and mouse models, DPY30 expression appears to be essential for the differentiation of embryonic stem cells as well as for hematopoietic, pancreatic, and neural progenitor/stem cells (38–44).

Here, in cancer cells, we identify a role for the WRAD core in maintaining genomic stability that is independent of its role in SET/MLL activity. We demonstrate that the WRAD core interacts with members of the DNA helicase complex, and, upon loss of DPY30, tumor cells undergo DNA re-replication, resulting in genomic instability. *In vivo*, loss of DPY30 increases the tumor mutational burden (TMB) and CIN, triggering a robust CD8-dependent anti-tumor immune response. Our findings implicate the WRAD core in preserving genome stability in cancer cells and suggest that DPY30 inhibition merits further exploration as a therapeutic concept to kill tumor cells by genome destabilization and sensitization to immune-checkpoint inhibitor therapy.

## Results

### Loss of DPY30 is dispensable for PDAC cell proliferation and H3K4 methylation

We first assessed the impact of silencing the expression of proteins comprising the WRAD core on PDAC cell proliferation. Consistent with our previous observations (31, 32), doxycycline-induced silencing of *WDR5* (*shWDR5*), *ASH2L* (*shASH2L*), or *RBBP5* (*shRBBP5*) strongly restrained cell proliferation *in vitro* (**Fig. 1a, Extended Data Fig. 1a-e**). Unexpectedly, shRNA-mediated silencing of *DPY30* (*shDPY30*) in human PDAC cells, as well as sgRNA-mediated *Dpy30* knockout (*Dpy30*-KO) in mouse PDAC cells from KPC (*Ptf1a^Cre^;KRas^lsl-G12D/wt^;Trp53^R172H/wt^)* (46) and KP^-/-^C (*Ptf1a^Cre^;KRas^lsl-fl^;Trp53^fl/fl^*) (47)mouse models did not affect cell proliferation (**Fig. 1b-c, Extended Data Fig. 1f-n**). We confirmed the lack of any effect of DPY30 loss on cell proliferation *in vivo*, showing that human *DPY30*-KO PDAC cells injected into immunodeficient NSG mice did not grow differently compared to controls (**Fig. 1d-g, Extended Data Fig. 1o-p**). Hence, our data show that PDAC cells are vulnerable to the loss of the WRA (WDR5, RBBP5, ASH2L) core, but not to DPY30 depletion.

**Figure 1.**
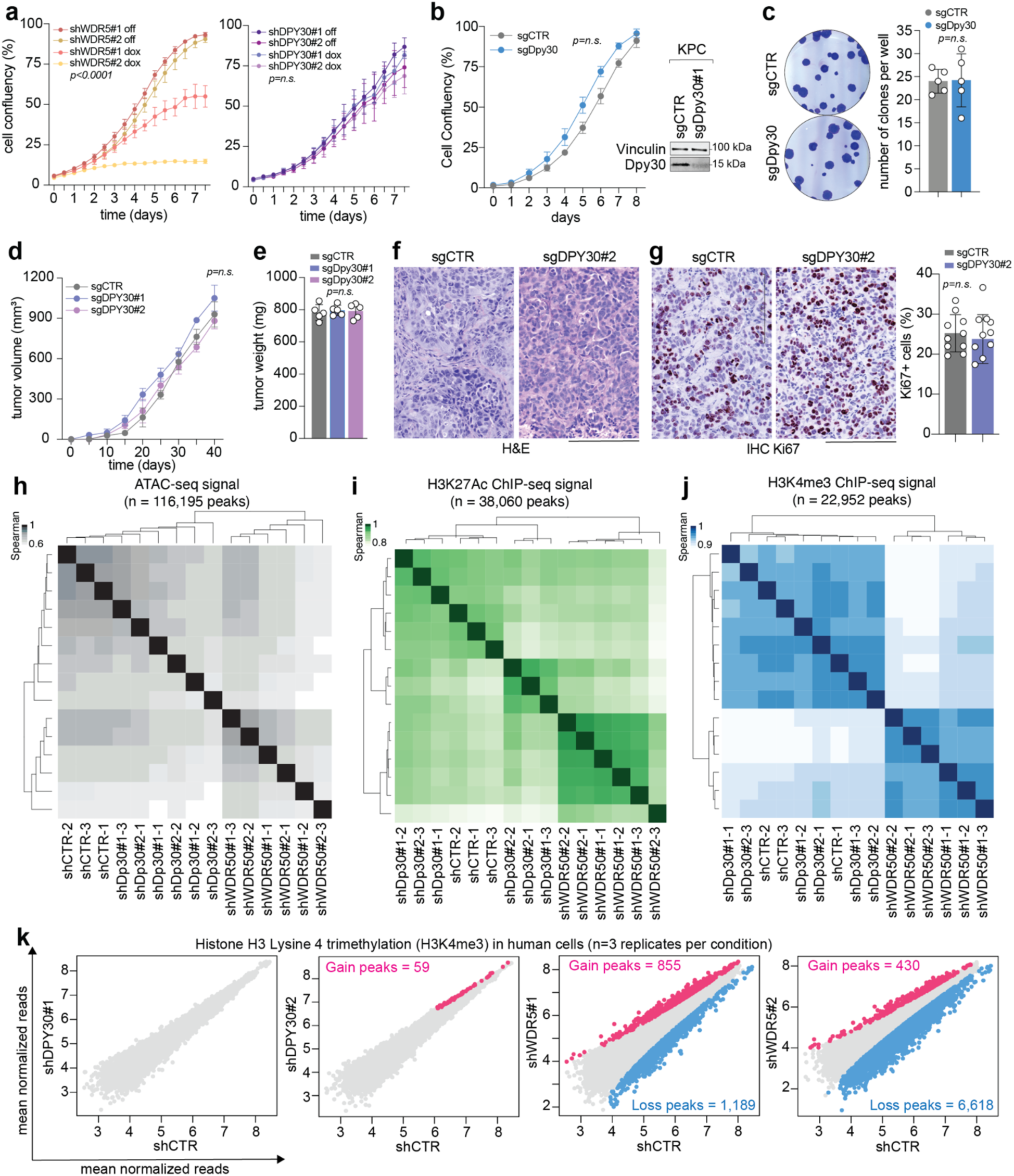
*DPY30* is dispensable for cancer cell proliferation and H3K4 trimethylation. **a.** Growth curve analysis of PATC53 cells carrying shRNAs targeting WDR5 (shWDR5), DPY30 (shDPY30) or control (shCTR) and treated with vehicle (off) or doxycycline (dox) to induce the gene silencing. **b.** Growth curve (left) and western blot (right) of control (sgCTR) or *Dpy30*-KO (sgDPY30) KPC cells. **c.** Clonogenic assay of KPC cells, as described in b, is shown (left) and quantified (right). **d-e.** Volume (d) and weight (e) of tumors from control (sgCTR) or DPY30-KO (sgDPY30) PATC53 cells grown in NSG mice. **f-g**. Micrographs showing the histology and Ki67 labeling in tumors described in (d). Hematoxylin and eosin (H&E) (f) staining, Ki67 IHC staining (g, left) and quantification (g, right) are shown. **h-j.** Heatmap showing Spearman correlations between samples and biological replicates of ATACseq (h), and H3K27ac (i) or H3K4me3 (j) Chip-seq in shCTR, shDPY30 or shWDR5 PATC53 cells. Samples are clustered by Euclidean distance and single-linkage clustering. **k.** Scatter plot showing mean normalized signals of H3K4me3 from samples described in j. Regions that significantly (FDR < 0.05) gained or lost H3K4me3 peaks in shDPY30 or shWDR5 versus shCTR are depicted in pink and blue, respectively. Not significant peaks are indicated as gray dots. In (a-e, g), data are expressed as mean ± SD, and dots represent biological replicates. Statistical significance was calculated using one-way ANOVA and Tukey’s multiple comparison tests (a, d, e), or Student t test (b, c, g). In panels f-g, scale bar = 200 μm.

Because the three-component WRA core has been shown to be the minimal required cofactor for supporting SET/MLL methyltransferase activity on histone H3K4, while the presence of DPY30 is only needed to enhance the specificity of the enzymatic activity (33), we explored whether inhibiting *DPY30* or *WDR5* expression in PDAC cells could differentially affect H3K4 methylation. To this end, we compared the acute effects of *DPY30* or *WDR5* loss on chromatin profiles in patient-derived PDAC cell lines transduced with doxycycline-inducible *shDPY30* or *shWDR5* by performing transposase-accessible chromatin using sequencing (ATAC-seq) and H3K4 tri-methylation (H3K4me3) and H3K27 acetylation (H3K27ac) chromatin immunoprecipitation sequencing (ChIP-seq), the latter of which provides readouts of COMPASS activity at gene regulatory regions (31, 48). Because WDR5 loss has a detrimental impact on the cell cycle (31, 39) (**Fig. 1a**), we excluded any confounding effects of WDR5 loss on proliferation by collecting cells at 48 hours after shRNA induction *via* doxycycline treatment, a timepoint sufficient for protein downregulation (**Extended Data Fig. 2a-b**). Cluster analysis of open genomic regions, as well as genomic regions marked by H3K4me3 and H3K27ac, revealed that *WDR5*-silenced cells exhibited chromatin states that were distinct from *DPY30*-silenced cells, which clustered with control cells. This suggests that, unlike the effects observed upon WDR5 loss, the acute and short depletion of *DPY30* is not sufficient to alter chromatin accessibility and states (**Fig. 1h-j**). Consistently, WDR5, but not DPY30, downregulation affected the total levels of H3K4me3 and H3K4me1 histone marks (**Extended Data Fig. 2b, f**).

Although genome-wide analysis of chromatin at regulatory regions proximal to gene transcriptional start sites (TSS) did not reveal significant differences between *DPY30* and *WDR5* silenced cells (**Extended Data Fig. 2c-e**), the H3K4me3 profiles of these two groups appear markedly different. Specifically, *DPY30* silencing did not affect the pattern of H3K4me3, indicating that DPY30 loss likely does not broadly alter the transcriptome of these cancer cells (**Fig. 1k**). In contrast, *WDR5* silencing significantly altered H3K4me3 patterns, resulting in the repression of thousands of genomic regions (**Fig. 1k, Extended Data Table S1**). To determine the functional annotations for the regions affected by WDR5 depletion, we conducted pathway analysis of genes located in these genomic regions and identified pathways involved in interferon alpha and gamma response, inflammatory response, and DNA damage (**Extended Data Fig. 2g, Extended Data Table S1**).

To dissect the biological role of the WRAD core in PDAC cells, we next performed bulk RNA-sequencing (RNAseq) analysis of patient-derived PDAC cells stably expressing *shDPY30*, *shWDR5*, or shCTR and treated with doxycycline for 48 hrs as described above (**Extended Data Fig. 2a**). *WDR5*, but not *DPY30*, silencing profoundly modulated the transcriptome of PDAC cells (**Extended Data Fig. 3a**), which is consistent with the essential role of WDR5 in regulating gene transcription (31, 36, 39, 49). Specifically, and in line with our epigenomics analysis (**Fig. 1k, Extended Data Table 1**), *WDR5* silencing induced a remarkable downmodulation of genes involved in the interferon alpha and gamma response as well as a dysregulation of pathways involved in mRNA translation and RNA processing (**Extended Data Fig. 3b, Extended Data Table S2**). Although *DPY30* silencing did not deregulate the transcriptome of PDAC cells as strongly as *WDR5* silencing, it induced a positive regulation of pathways involved in interferon signaling, inflammatory response, and DNA damage (**Extended Data Fig. 3e-g, Extended Data Table 2**). Overall, our data indicate that *DPY30* or *WDR5* silencing leads to distinct epigenomic and transcriptomic changes in PDAC, thus demonstrating that the WRAD core has non-redundant roles in maintaining cell proliferation and transcriptional activity in PDAC.

### WRAD core directly interacts with the MCM complex

To further investigate the discrepancies in the proliferative, epigenetic, and transcriptional phenotypes that emerged upon depletion of different WRAD core members in PDAC cells, we expanded our investigation into the biological function of DPY30 in PDAC. To this end, we immunoprecipitated endogenous DPY30 protein in two human (PATC53 and PATC66) and one mouse (KPC) PDAC-derived cell models, all of which carry *TP53* and *KRAS* mutations (31, 32). Unbiased proteomic analysis showed that multiple top-scoring interactors of DPY30 are involved in chromatin remodeling, RNA metabolism, splicing, and gene transcription (**Extended Data Fig. 4a-d, Extended Data Table S3**), consistent with the role of DPY30 as a cofactor supporting histone methylation and gene transcription regulation by COMPASS complexes (38–44). Proteins involved in DNA replication, DNA damage, and cell cycle regulation were also identified as DPY30 interactors (**Extended Data Fig. 4a-d, Extended Data Table S3**). These included proteins previously unknown to interact with DPY30, such as multiple members of the DNA replisome, including the MCM complex and WDHD1 (**Fig. 2a, Extended Data Fig. 4d**). We validated these findings by expressing either mouse or human HA-tagged DPY30 with mouse or human MCM subunits and found that DPY30 bound most strongly to MCM6 compared to others (**Fig. 2b, Extended Data Fig. 4e**). Notably, proximity ligation analysis (PLA) revealed that endogenous DPY30 lies in proximity (<40nm) of both MCM6 and WDHD1, strongly suggesting that DPY30 may be a core component of the DNA replisome (**Fig. 2c)**. To explore whether the interaction of DPY30 with the replisome was cell cycle-dependent, we expressed the fluorescent ubiquitination-based cell cycle indicator (FUCCI) system in human PDAC cells (50). By enabling the real-time tracking of cell cycle progression in live cells (54), this approach allowed us to sort PDAC cells by cell cycle phase and then perform immunoprecipitation (IP) assays for endogenous DPY30 (**Extended Data Fig. 4f-g**). Unbiased proteomic analysis of affinity-purified samples showed that members of the COMPASS complex were, as expected, enriched in DPY30 pull-downs and that these interactions were independent of cell cycle phase (**Extended Data Fig. 4h**). Furthermore, this analysis confirmed that WDHD1 and all MCM complex members, including MCM10, MCM3AP, and MCMBP, interacted with DPY30, with all replisome components except for MCM2, MCM4, and MCM10 interacting with DPY30 in a cell cycle-independent fashion **(Extended Data Fig. 4i**). Additionally, we found that WDR5 also interacted with MCM complex members (**Fig. 2d)**. In cell nuclei, WDR5 was shown to reside topographically close to MCM6 and WDHD1 (**Fig. 2e-f, Extended Data Fig. 4j**), similar to what was seen for DPY30 (**Fig 2c**). Importantly, endogenous WDR5, ASH2L, and RBBP5 could be precipitated with MCM6, suggesting that the whole WRAD participates in this interaction (**Extended Data Fig. 4k-m**). Interestingly, destabilization of the WRAD core through silencing of either *WDR5* or *DPY30* impeded the endogenous binding of DPY30 to MCM6 (**Fig. 2g-h).**

**Figure 2.**
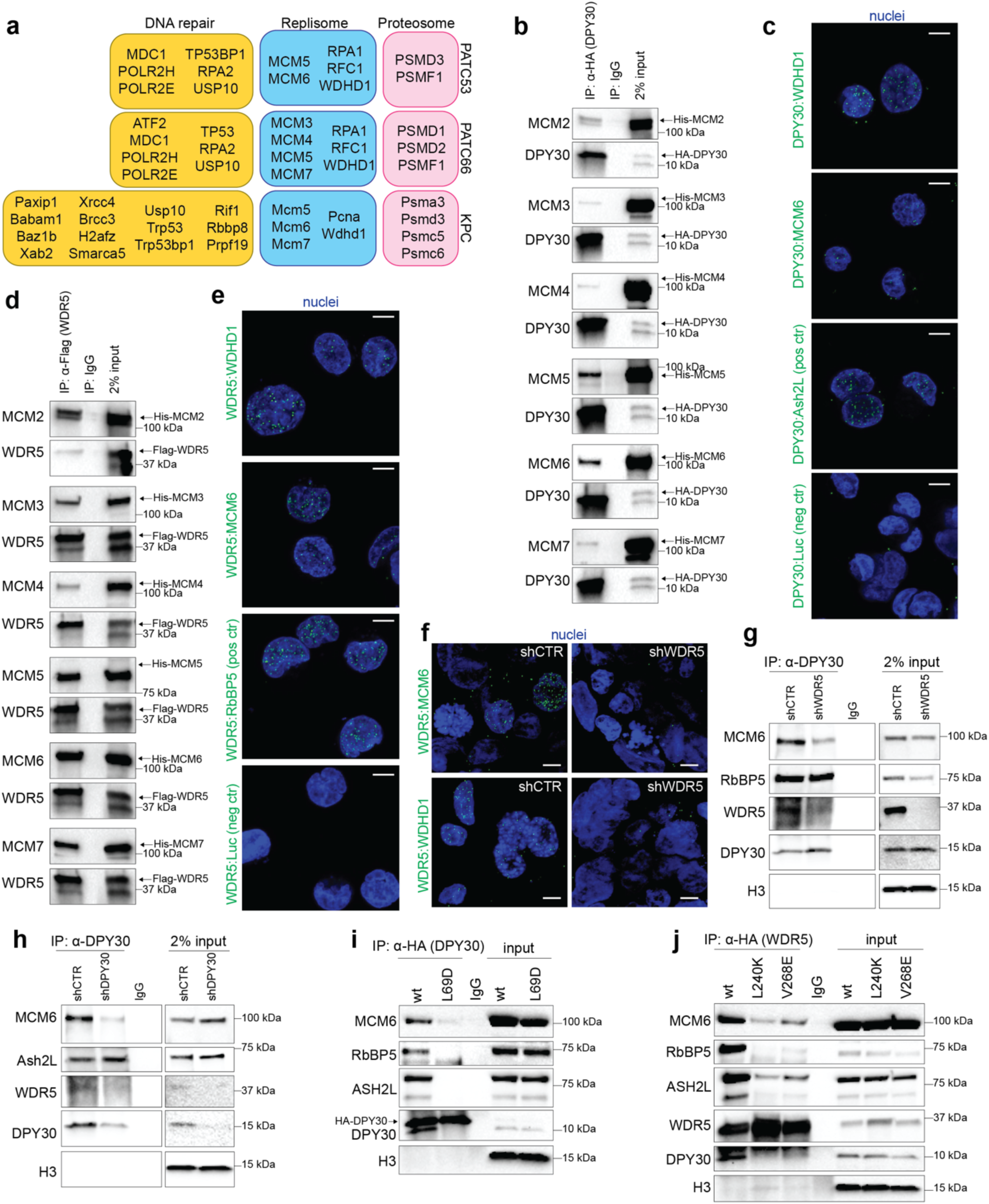
The WRAD core interacts with WDHD1 and the MCM complex. **a.** Schematic representation of DPY30 interactors involved in DNA repair, replisome, and proteosome signaling pathways in PATC53, PATC66, and KPC cells. **b.** Western blot analysis of the indicated proteins in 293T cells co-overexpressing HA-tagged human *DPY30* and His-tagged human MCM complex subunits. Control IgG and input are shown. **c.** Proximity Ligation Assay (PLA) staining detecting DPY30 interaction with WDHD1, MCM6, Ash2L (positive control) or Luciferase (negative control). **d.** Western blot analysis of the indicated proteins in 293T cells co-overexpressing Flag-tagged human *WDR5* and His-tagged human MCM complex subunits. Control IgG and input are shown. **e.** PLA staining detecting WDR5 interaction with WDHD1, MCM6, RbBP5 (positive control) or Luciferase (negative control). **f.** PLA staining detecting WDR5 interaction with WDHD1 and MCM6 in PATC53 control (shCTR) or silenced for WDR5 (shWDR5). **g-h.** Western blot analysis of the indicated proteins in PATC53 cells control (shCTR) or silenced for WDR5 (shWDR5, g) or DPY30 (shDPy30, h). Control IgG and input are shown, and H3 was used as a loading control. **i-j.** Western blot analyses of the indicated proteins in KPC cells control or overexpressing DPY30 wild type (WT) or mutant (L69D) isoforms (i), and in KPC cells control or overexpressing WDR5 wild type (WT) or mutant (L240K, V268E) isoforms (j). Control IgG and input are shown, and H3 was used as a loading control. In figure, scale bar = 5 μm.

To explore whether WRAD core integrity is necessary for MCM6 binding, we overexpressed a mutant isoform of DPY30 (L69D) known to hinder the DPY30:ASH2L interaction and the proper assembly of the WRAD core (51). Remarkably, DPY30^L69D^ expression prevented the DPY30:MCM6 interaction as well as the assembly of the WRAD core **(Fig. 2i, Extended Data Fig. 4n**). We employed a similar approach by overexpressing two WDR5 mutant isoforms (L240K and V268E) that have altered WDR5-binding motif (WBM) domain and are known to disrupt the interaction of WDR5 with RBBP5 and cMyc (31). We consistently found that disrupting the WRAD core assembly strongly reduced the interaction between WDR5 and MCM6 **(Fig. 2j, Extended Data Fig. 4o**). Altogether these observations suggest that the stability of the WRAD core is necessary for the interaction between the WRAD core and the MCM6 subunit of the MCM complex.

### The WRAD core participates in replication fork dynamics

Because the MCM complex represents the DNA helicase activated at the start of DNA replication (10, 12–15, 23), we tested whether WRAD contributes to replication fork progression. Using an iPOND assay to recover proteins bound to newly synthesized DNA, we found that WRAD core members localized at DNA replication fork initiation sites (2 min of EdU pulse), persisted during DNA elongation (5-10 min of EdU pulse), but disappeared at later time points (chase) **(Fig. 3a, Extended Data Fig. 5a**). Notably, histone H3, a marker for nucleosome incorporation and an indicator of SET/MLL canonical activity, was barely detectable at 2 min of EdU pulse, gradually incorporated at 5 and 10 minutes, and strongly detected during the chase (**Fig. 3a, Extended Data Fig. 5a**). This contrasting behavior between WRAD and H3 serves as an internal control, confirming that WRAD members detected at DNA replication forks are not associated with chromatin-bound proteins (**Fig. 3a, Extended Data Fig. 5a). These data strongly indicate that, in the context under study, the WRAD core directly interacts with the DNA replication fork in a nucleosome-independent manner. Importantly, analysis of the interactions between proteins and the nascent DNA using a single cell-based method (SIRF assay) (52, 53**) showed that WDR5, but not DPY30, directly interacts with nascent DNA, further supporting a pivotal role for the WRAD core in DNA replication dynamics (**Fig. 3b**).

**Figure 3.**
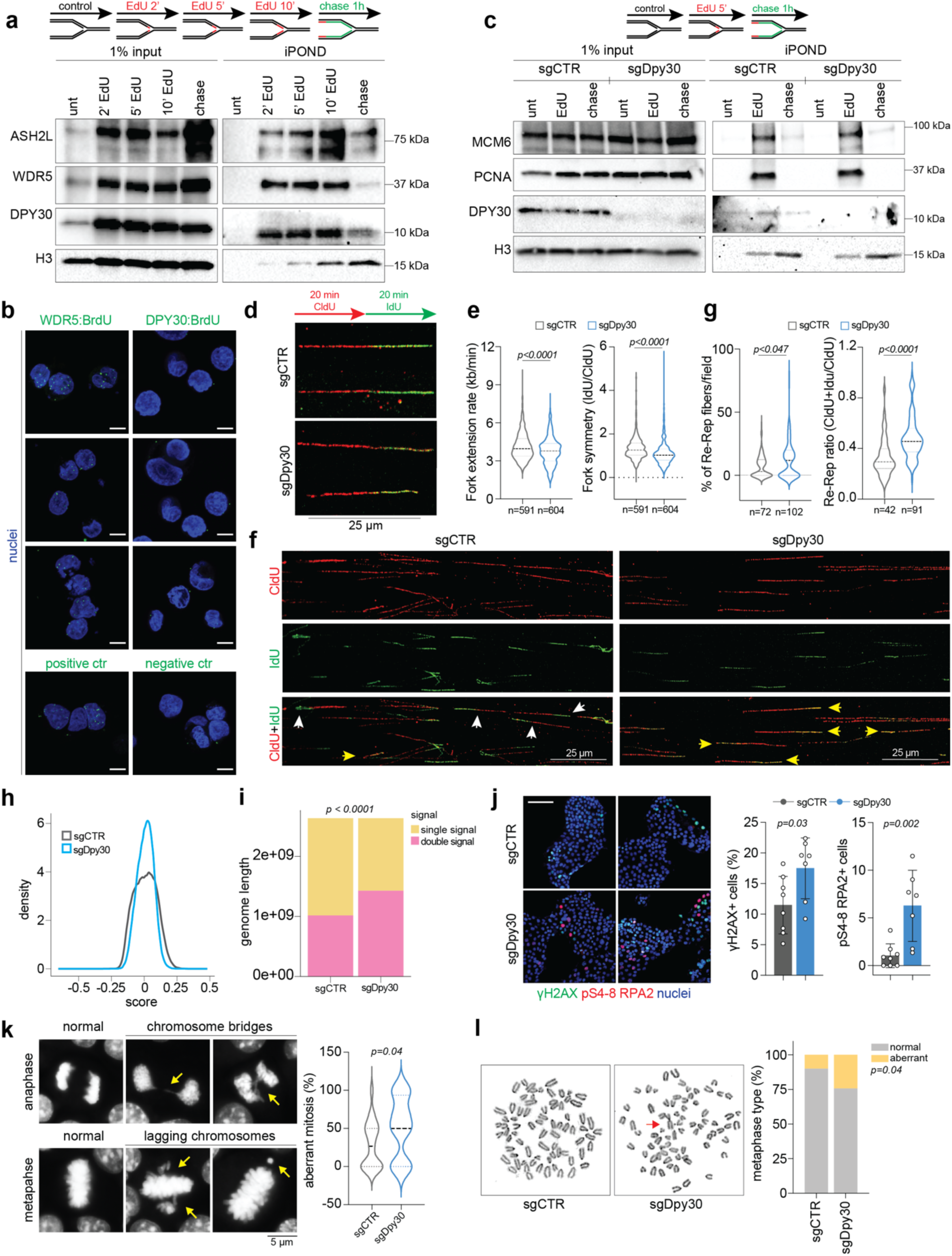
DPY30 loss leads to DNA re-replication and CIN. **a.** Western blot analysis of proteins bound to nascent DNA retrieved using iPOND or input control in KPC cells. Cells were treated as follows: control (negative control, no biotin in the click reaction), EdU pulse (EdU treatment for 2, 5 or 10 min), chase (EdU treatment for 10 min, followed by a 60 min thymidine chase). Histone H3 was used as a loading control. **b.** Modified PLA (SIRF) assay detecting the interaction between the incorporated BrdU (5 min pulse) and WDR5 (left) or DPY30 (right). BrdU:PCNA and BrdU:luciferase were used as positive and negative control, respectively. Scale bar = 5 μm. **c.** Western blot analysis of iPOND or input control in control (sgCTR) or *Dpy30*-KO (sgDpy30) KPC cells. Cells were treated as described in a. Histone H3 was used as a loading control. **d.** Schematic illustration and confocal images of DNA fiber assay in sgCTR and sgDpy30 KPC cells. **e.** Violin plots reporting mean fork progression rate (left) and fork symmetry (right) in cells as in d. **f.** Confocal images of DNA fiber assay in sgCTR or sgDpy30 KPC cells. White arrows indicate a fiber with non-aberrant DNA replication, yellow arrows indicate overlapped CldU and IdU incorporation (DNA re-replication). **g.** Violin plots reporting the frequency of DNA re-replication (left) and the fiber re-replication ratio (right) in cells as in f. **h.** Trend curves showing the dual CldU:IdU incorporation (score) calculated from the density of ChIPseq reads of IdU and CldU IP in sgCTR and sgDpy30 KPC cells. **h.** Bar plot showing genome length of single (IdU or CldU) and double nucleotide (CldU+IdU) incorporation in sgCTR and sgDpy30. **j.** immunofluorescence images of sgCTR or sgDpy30 KPC cells stained for pS4-8 RPA (red), pS139 H2AX (γH2AX, green), and nuclei (blue). Right, graphs reporting the percentage of γH2AX-positive or pS4-8 RPA-positive sgCTR or sgDpy30 KPC cells. Each dot represents a different analyzed field, scale bar = 100 μm. **k.** Left, immunofluorescence images of mitotic KPC cells stained with DAPI (DNA, white). Yellow arrows indicate lagging chromosomes and chromosome bridges. Right, violin plot reporting the mean percentage of sgCTR or sgDpy30 KPC cells with aberrant mitotic outcomes *in vitro*. **k.** Left panel, images of metaphase spreads in sgCTR or sgDpy30 KPC cells stained with Giemsa (chromosome). Red arrow indicates chromosome fusion and fragmentation. Right panel, percentage of KPC cells with normal (gray) or aberrant (yellow) mitotic outcomes. In the figure, n indicates the number of analyzed events. Data are expressed as mean values ± SD. Statistical significance was calculated using the Mann-Whitney test (e, g), Student’s *t* test (j, k) or Fisher’s exact Chi square test (i, l).

Because WDR5 deregulation affects distinct biochemical pathways and results in rapid induction of cell death (**Fig. 1**) (31), we sought to avoid any confounding effects by focusing further mechanistic studies on DPY30. First, iPOND analysis found that *Dpy30*-KO did not destabilize MCM6 nor WDR5 binding at replication forks, suggesting that DPY30 is not directly involved in regulating fork assembly (**Fig. 3c, Extended Data Fig. 5b**). Second, to examine the effect of DPY30 on replication fork efficiency, we performed a DNA fiber assay in *Dpy30*-KO and control cells. *Dpy30*-KO resulted in slower fork progression rates and reduced symmetry between incorporation of the two different nucleotide analogues, suggesting that *Dpy30* loss may promote replication stress (**Fig. 3d-e**). Additionally, we observed increased frequency of overlap between the two incorporated nucleotide analogues in *Dpy30*-KO cells compared to controls, indicating DNA re-replication in these cells, and further supporting the hypothesis that *Dpy30* loss may contribute to replication stress (**Fig. 3f-g**).

To corroborate our findings, we next developed an unbiased DNA re-replication sequencing assay (“Re-repli-seq”, see methods section) (Extended Data Fig. 5c). Specifically, S phase synchronized KPC controls or Dpy30-KO cells were released in presence of CldU (45 min) followed by a second pulse of IdU (45 min) (Extended Data Fig. 5c). The cells were then split, and a ChIP assay was performed using antibodies against the incorporated CldU and IdU separately, followed by next generation sequencing. After, sequence alignment to reference genome, we represented the data as a ratio of incorporated CldU:IdU (38). In case of DNA re-replication, sequencing of double-labelled immunoprecipitated genomic regions should result in an increased overlap between CldU and IdU sequencing reads. Data analysis showed that, compared to controls, the Dpy30-depleted KPC cells showed broader genomic domains with CldU:IdU ratio close to zero, which means the nucleotide analogues are incorporated equally and overlap in the same genomic regions (**Figure 3h-i**). As an example, regions of different chromosomes (Chr1, Chr2, Chr4, Chr6, Chr10 and Chr16) displayed domains of replicated DNA with an overlapped pattern of CldU:IdU ratio more prominently distributed in Dpy30-depleted cells compared to control cells, that instead, showed a more coordinated replication profile (Extended Data Fig 5d-e). Our genome-wide analysis of CldU:IdU incorporation strongly supports that Dpy30 and the WRAD core are required to sustain a coordinated DNA replication during the S phase in tumor cells.

Because excessive DNA replication can trigger DNA damage, we found that during unperturbed growth condition, *Dpy30*-KO KPC cells showed stronger γH2AX (pS139) and pRPA (pS4-8) staining compared to control cells (**Fig. 3j, Extended Data Fig. 5f-h**). Consistently, γH2AX levels *in vivo* were dramatically increased in both *Dpy30*-KO KPC and human-derived *DPY30*-KO or knockdown (KD) tumors compared to controls (Extended Data Fig. 6a-b). Further, bulk RNAseq analyses performed in mouse-derived Dpy30-KO and control PDAC cells revealed a deregulation of pathways involved in DNA damage and G2/M checkpoint (Extended Data Fig. 6c-d, Extended Data Table S4), Interestingly, Dpy30-depleted cells showed an extensive upregulation of inflammatory pathways (Extended Data Fig. 6c-d, **Extended Data Table S4)**, which might have been activated as a consequence of persistent DNA damage (2, 55, 56). These data align with our previous work (31) showing that the WDR5 silencing strongly increased DNA damage and induced a sustained G2/M accumulation in PDAC cells. Further, these observations suggest that PDAC cells can tolerate the incremental DNA damage caused by *DPY30* loss, but cannot survive the extensive DNA injuries and transcriptomic rewiring upon WDR5 depletion. This suggests that the intact WRAD core sustains genomic stability by counteracting excessive DNA replication.

### DPY30 maintains genome integrity by limiting excessive DNA replication and stabilizing the S-to-G2/M transition

The risk of DNA re-replication increases at the S-to-G2/M boundary, during which Cdc6 and Cdt1 are in their active forms and the expression of geminin, an inhibitor of DNA replication, has not yet peaked (10, 13, 23). To investigate whether *DPY30* loss disrupts cell cycle dynamics in these phases specifically, we synchronized PDAC cells in S phase using a double-thymidine block and collected cells at different time points after release (Extended Data Fig. 6e-f). In line with our observations in proliferation assays (**Fig. 1a-d, Extended Data Fig. 1a-n**), we observed that *Dpy30* KO did not grossly affect cell cycle distribution in non-synchronized cells (Extended Data Fig. 6e-f). However, in synchronized cells, a larger proportion of *Dpy30*-KO cells compared to control cells advanced towards the G2/M boundary by 2 hours after release from S phase (Extended Data Fig. 6e-f). Additionally, at this time point, we detected an increased percentage of aneuploid cells (DNA content >4N), higher DNA damage, increased BrdU incorporation, as well as higher phosphorylation of pS139 MCM2, pS54 Cdc6, and pS10 histone H3 in Dpy30-KO cells compared to controls, all of which support an uncoupling of DNA replication and cell cycle progression upon *Dpy30* loss (Extended Data Fig. 6e-h). We further observed that, while there were no significant differences in the expression of geminin in the arrested cells (Extended Data Fig. 6h-i), geminin expression was strongly downmodulated upon release in *Dpy30*-KO cells. We next synchronized *Dpy30*-KO or control PDAC cells in S phase using a double thymidine block, and in G2/M phase using a thymidine-nocodazole block (Extended Data Fig. 6j). Strikingly, Western blot analysis showed that the dysregulation of Cdc6 and geminin expression was restricted to the G2/M phase rather than the early S phase (Extended Data Fig. 6j). Overall, our data indicate that, upon DPY30 depletion, the risk of DNA re-replication increases at the S-to-G2/M boundary due to the uncoupled regulation of the Cdc6/Geminin axis (20, 24, 57–59).

During the G2/M and G1 phases, Geminin stability is tightly regulated by the anaphase promoting complex/cyclosome (APC/C), leading to its proteasome-dependent degradation (60). As a member of the E3 ubiquitin ligase family, APC/C requires the modification of the cullin ring with NEDD8, an-ubiquitin-like protein, for its activity (61).

Chemical inhibition of NEDD8 promotes DNA re-replication through the stabilization of Cdt1 and Cdc6 protein expression (24, 61–63). Hence, we hypothesized that NEDD8 inhibitors, such as MLN4924, may exacerbate the uncoupling between DNA replication and G2/M transition in Dpy30-depleted cells by. As expected, human nearly diploid RPE-1 cells treated with MLN4924 showed increased expression of Cdc6 in both control and *DPY30*-KO cells compared to untreated ones (Extended Data Fig. 7a). Further, when compared to control RPE-1 cells, geminin expression was remarkably lower upon DPY30 depletion and MLN4924 treatment, and the spontaneous activation of γH2AX was observed upon DPY30 depletion and then boosted by MLN4924 treatment (Extended Data Fig. 7a). Finally, we observed increased BrdU incorporation in cells treated with MLN4924, with the highest incorporation in *DPY30*-KO cells (Extended Data Fig. 7b). These data were confirmed in KPC cells, showing dysregulated geminin and γH2AX expression, as well as increased percentage of aneuploid cells, in *Dpy30*-KO compared to control cells (Extended Data Fig. 7c-d). Overall, our data suggested that *Dpy30* loss leads to DNA re-replication through a potent perturbation of the Cdc6/geminin balance during the G2/M phase.

Finally, because uncoordinated replication can contribute to chromosomal instability (CIN), we investigated whether DPY30 may induce this phenomenon (16, 22, 25, 64). Indeed, *DPY30*-KO in both human and mouse PDAC-derived models increased the percentage of cells with chromosome mis-segregation and micronuclei *in vitro* and *in vivo (***Fig. 3k, Extended Data Fig. 7e-g**). Consistently, metaphase spreads showed an increased in the number of chromosomal breaks and fusions *in vitro* (**Fig. 3l, Extended Data Fig. 7h-i**). Collectively, these data support a critical role for DPY30 in maintaining genome stability and suppressing DNA re-replication at the G2/M transition.

### *DPY30*-KO cells display complex karyotypes and elicit a T cell-mediated immune response

To assess whether the genome instability induced by DPY30 loss may impact tumor immunogenicity *in vivo*, we injected control or *Dpy30*-KO KPC and KP^-/-^C cells into cohorts of NSG, nude, and C57BL/6 mice. Strikingly, *Dpy30*-KO strongly impaired tumor growth compared to controls in immune-competent C57BL/6 mice but had no effect on tumor growth in immunodeficient mice (**Fig. 4a-d, Extended Data Fig. 8a-d**). Moreover, histological analysis revealed that *Dpy30*-KO tumors grown in C57BL/6 mice appeared more differentiated and characterized by an increased number of mucin-positive ductal-like structures compared to their respective controls (**Fig. 4e-f**). Of note, in models of orthotopic transplantation, we observed that Dpy30 depletion significantly prolonged animal lifespan (**Fig. 4g, Extended Data Fig. 8e**).

**Figure 4.**
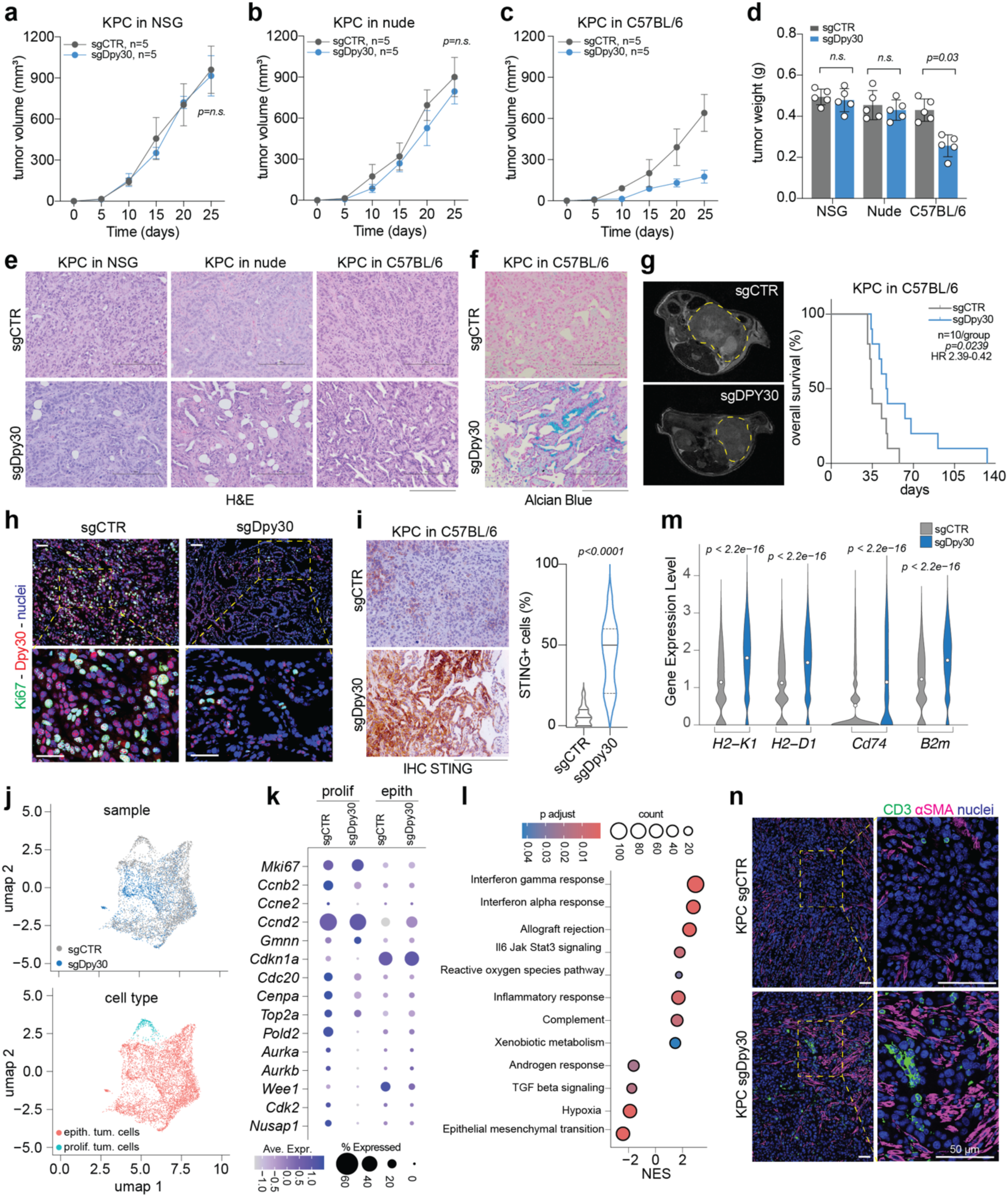
DPY30 loss results in a distinct immune context-dependent phenotype **a**−**c.** Volume of control (sgCTR) and *Dpy30*-KO (sgDpy30) KPC tumors grown in the flank of immune-deficient NSG (a), nude (b), or in immune-competent C57BL/6 mice (c). **d.** Weight (mg) of tumors described in (a−c). **e.** Hematoxylin and eosin (H&E) staining of tumors from experiments described in (a−d). **f.** Alcian blue staining of tumors from sgCTR and sgDpy30 KPC cells grown in C57BL/6 mice. **g.** Left, T2 Magnetic Resonance Imaging (MRI) scans of tumors from sgCTR and sgDpy30 KPC cells grown orthotopically in the pancreas of C57BL/6 mice. Right, Kaplan-Meier survival curve of tumor-bearing mice. Statistical significance and Hazard Ratio (HR) were calculated using the Log-rank test. **h.** Immunofluorescence staining of Ki67 and DPY30 in sgCTR and sgDpy30 KPC cells grown in C57BL/6 mice. **i.** Left, STING immunohistochemical (IHC) staining of tumors from sgCTR and sgDpy30 KPC cells grown in C57BL/6 mice Right, violin plot showing the percentage of STING positive cells per field. **j.** UMAP plots showing bi-dimensional cluster distribution of sgCTR (n=3) or sgDpy30 (n=3) KPC tumor epithelial cells after filtering and quality control. **k.** Bubble plot showing expression levels of the genes involved in cell cycle analyzed from the single-cell RNA sequencing data as described in j. Size of dots represents the percentage of cells expressing the gene; color scale shows the average expression level **l.** Bubble plot showing the over-representation analysis of genes enriched in sgDpy30 KPC cells. Gene expression was calculated using Seurat. **m.** Violin plot showing the expression of *B2m, H2-K1, H2-D1 and Cd74* per cell calculated using Seurat in sgCTR and sgDpy30 KPC cells. **n.** Immunofluorescence staining of CD3 and αSMA in sgCTR or sgDpy30 KPC cells grown in C57BL/6 mice. In the figure, n indicates the number of mice analyzed. In (a−d), data are expressed as mean ± SD and statistical significance was calculated using an unpaired One-way ANOVA (a-c) or Student’s *t* test (d). In (I, m), data are expressed as median values and statistical significance was calculated using the Mann Whitney test. In (e−f, i), scale bar = 200 μm. In (h, n), scale bar = 50 μm.

To eliminate potential confounding effects due to CRISPR/Cas9-related immunogenicity (65), we injected mouse cells derived from KP^-/-^C tumors stably expressing doxycycline-inducible shDpy30 or shCTR into NSG or C57BL/6 mouse backgrounds. Also in this setting, we confirmed that Dpy30-depleted tumors grown in immunocompetent mice were smaller and more differentiated compared to controls, while in NSG mice no major differences were detected (Extended Data Fig. 8f-i). Further, we confirmed that *Dpy30* silencing strongly prolonged animal lifespan (Extended Data Fig. 8j).

Intriguingly, the cell proliferative index, evaluated as Ki67- and pS10-H3-positive staining, was remarkably reduced in *Dpy30*-KO cells compared to controls in immune-competent, but not NSG, backgrounds (**Figure 4h, Extended Data Fig. 5g** and 8k-l). This suggests that, in the absence of DPY30, the immune system may inhibit growth of or preferentially clear proliferating tumor cells. Interestingly, gene set enrichment analysis (GSEA) from bulk RNAseq data of *Dpy30*-KO and control KP^-/-^C tumors showed a positive enrichment score for pathways involved in DNA damage and repair process in immunodeficient mice, while the same cancer cells showed upregulation of gene pathways involved in inflammation, including interferon signaling, the complement cascade, and PD-1 signaling, in an immunocompetent background (Extended Data Fig. 8m-n**, Extended Data Table S5**). Because CIN can enhance the activity of the cGAS/STING pathway, which in turn regulates the expression of the type I interferons and related gene products (56), we sought to evaluate the expression of STING upon Dpy30 depletion. Immunostaining analysis of KPC showed a potent activation of STING in *Dpy30*-KO cells compared to controls both *in vitro* and *in vivo* **(Fig. 4i, Extended Data Fig. 8o**). Altogether, these data suggest that *Dpy30* loss can trigger a T cell-mediated immune response *in vivo* through DNA damage and CIN-dependent activation of the type I interferons, the latter of which occurs through the cGAS/STING pathway.

Next, we performed single-cell RNAseq (scRNAseq) on *Dpy30*-KO and control KPC tumors (n=3 per condition) grown in immunocompetent mice. Dimensionality reduction analysis of sub-clusters of tumor cells revealed only a partial overlap between the transcriptomes of *Dpy30*-KO and control tumor cells (**Fig. 4j)**. Specifically, we identified a population of highly proliferative tumor cells that was expanded in control tumors but minimally represented in *Dpy30*-KO tumors (**Fig. 4j-k**). Expression analysis of cell cycle regulators corroborated our observation, showing a reduced expression of proliferative genes (*Ccnb2, Cdc20, Top2a, Aurka, Aurkb, Wee1*) (**Fig. 4k)** and, consequently, supporting the hypothesis that highly proliferative tumor cells are selectively cleared by the immune response in the absence of Dpy30. Further, GSEA confirmed an upregulation of genes involved in interferon signaling and inflammatory response pathways (Il6/Stat3, allograft rejection, and complement) upon Dpy30 depletion (**Fig. 4l**). Remarkably, *B2m*, *Cd74*, *H2-K1*, and *H2-D1*, genes involved in antigen presentation, were among the top-scoring genes in *Dpy30*-KO cells (**Fig. 4m**). Indeed, when comparing *Dpy30*-KO tumors to controls, immunostaining analysis confirmed an increased infiltration of CD3+ tumor-infiltrating lymphocytes (TILs) as well as an increase of cancer associated-fibroblasts (CAFs) (αSMA+), the latter likely the result of the depletion of the epithelial compartment (**Fig. 4n, Extended Data Fig. 8p**). Accordingly, examination of the tumor microenvironment (TME) at the single cell level revealed an increased number of CAFs, B cells, NK cells, and T cells in *Dpy30*-KO tumors grown in an immunocompetent background when compared to controls (**Fig. 5a-c, Extended Data Fig. 9a-b**). These findings were corroborated by flow cytometry-based immunophenotyping, which demonstrated a significant increase in the number of all infiltrating T cell subsets and NK cells in *Dpy30*-KO tumors. Among the immune cells, the CD8+ T cells were the predominant population, suggesting that they may be the primary contributor to the anti-tumor response (**Fig. 5d, Extended Data Fig. 9c-e**). Importantly, compared to control tumors, CD8+ T cells in *Dpy30*-KO tumors displayed lower expression of exhaustion marker *Tox* as well as higher expression of cytotoxic markers such as interferon gamma (*Ifng*), granzyme B (*Gzmb*), and natural killer cell granule protein 7 (*Nkg7*) (**Fig. 5e, Extended Data Fig. 9f-h**). Altogether, these findings suggest that CD8+ T cells are activated and may selectively clear *Dpy30*-KO tumor cells with increased immunogenicity due to genomic instability.

**Figure 5.**
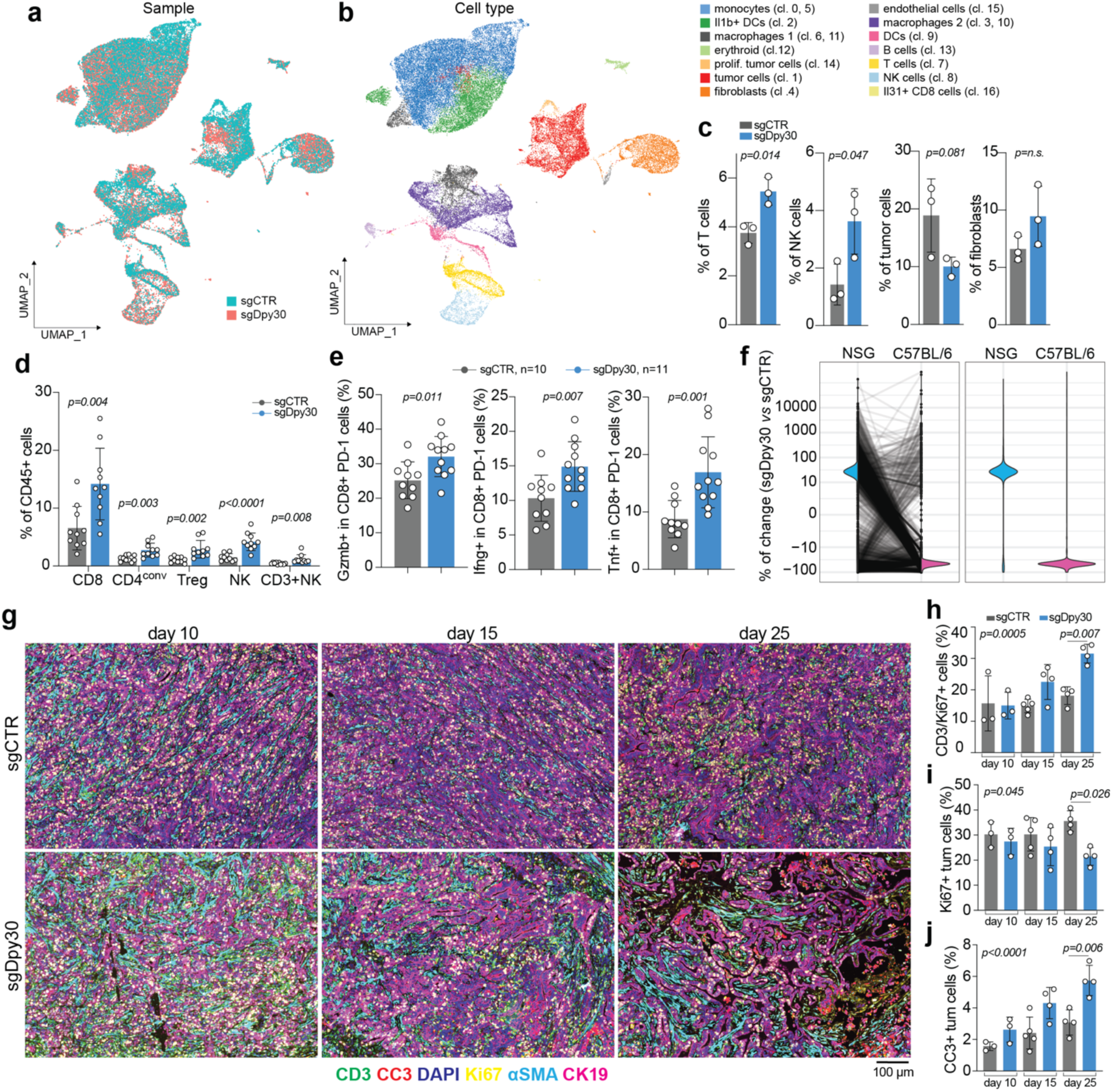
*Dpy30*-KO tumors display higher T cell infiltration, activation, and heterogeneity. **a-b.** UMAP plot shows bi-dimensional sample (a) and cell type (b) distribution of control (sgCTR) or *Dpy30*-KO (sgDpy30) KPC tumors after filtering and quality control calculated using Seurat. **c.** Cell fraction of T cells, NK cells, tumor cells and fibroblasts calculated using Seurat in sgCTR and sgDpy30 KPC tumors. **d.** Percentage of CD8+ T cells, CD4+ conventional T cells, CD4+/Foxp3+ T regulatory cells, CD3+ NK, and NK cells in sgCTR or sgDpy30 KPC tumors grown in immune-competent C57BL/6 mice. Data were analyzed using flow cytometry. **e.** Percentage of CD8+/PD-1+ T cells in sgCTR or sgDpy30 KPC tumors grown in C57BL/6 mice and expressing the cytotoxic markers *Gzmb, Ifng, Tnf* as analyzed by flow cytometry. **f.** Violin plots showing the percentage of barcode changes in sgCTR and sgDpy30 KPC injected in NSG (blue) or C57/BL6 (pink) background. **g.** Multiplexed immunofluorescence staining of CD3 (green), cleaved caspase 3 (CC3, red), Ki67 (yellow), αSMA (cyan), cytokeratin 19 (CK19, magenta) and DAPI (blue) in sgCTR or sgDpy30 KPC cells grown in C57BL/6 mice and collected after 10, 15 or 25 days after cell injection. **h-j.** Bar plot showing the percentage of CD3/Ki67+ cells (h), CC3+ cells (i) or Ki67+ cells (j) in the samples described in g. In (c-e, h-j) each dot represents a biological replicate and data are presented as mean ± SD and statistical significance was calculated using Student’s *t* test (c-e) or one-way ANOVA (h-j).

To test this hypothesis, whole-exome sequencing (WES) was performed on *Dpy30*-KO or control tumors grown in NSG or C57BL/6 mice. We found that, in a permissive immune-deficient environment, *Dpy30*-KO tumors grew without restraint and tolerated a higher mutational burden (TMB) compared to *Dpy30*-proficient controls (Extended Data Fig. 9i). However, the same cells transplanted in a selective immune-competent background displayed lower TMB (Extended Data Fig. 9i), thus suggesting that cancer cells undergoing uncoordinated DNA replication and CIN were being cleared from tumors. To corroborate our findings and quantify the extent of this depletion, we generated barcoded KPC cells using our established platform (66, 67). We then transiently infected barcoded KPC cells with sgRNAs targeting *Dpy30* or a scramble control and injected them into NSG or C57BL/6 mice. Explanted tumors were subjected to next-generation sequencing (NGS) for barcode detection. Pairwise comparison of subclonal composition of *Dpy30*-depleted versus control KPC cells grown in immunodeficient or immunocompetent mice showed a dramatic depletion of barcodes in the *Dpy30*-KO cells transplanted in C57BL/6 mice compared to an immunodeficient counterpart (**Fig. 5f**). Further, we injected *Dpy30*-KO or control KPC cells into immunocompetent C57BL/6 mice and collected tumors at different time points (day 10, day 15 and day 25). Multiplexed immunostaining analysis showed that, over time, the number of TILs strongly increased exclusively in *Dpy30*-KO tumors compared to controls (**Fig. 5g-h**). This immune infiltration, in the *Dpy30*-KO tumors, was paralleled by a net increase in the cleaved caspase 3 (CC3) staining, a *bone fide* marker of apoptotic cell death, as well as a pronounced reduction in the number of Ki67+ tumor cells (**Fig. 5i-j**). Altogether these data indicate that, in a mutated *TP53/KRAS* genetic background, increases in CIN and TMB due to DPY30 loss enhance the immunogenicity of PDAC cells and a consequent T cell-mediated response.

### DPY30 expression correlates with poor response to immune checkpoint blockade and tumor staging

To determine whether CD8+ T cells are responsible for clearing cells harboring a high TMB (high-TMB) due to DPY30 depletion, we injected *Dpy30*-KO or control KPC cells into C57BL/6 mice and depleted the CD8+ T cell population using an anti-CD8 monoclonal antibody (**Fig. 6a**). We found that CD8+ T cell depletion was sufficient to rescue the phenotype induced by *Dpy30* loss, considerably accelerating tumor growth, and reducing lifespan when compared to that of control mice (**Fig. 6b-c**). To explore whether the activation of cytotoxic T cells upon *Dpy30* loss may synergize with immunotherapy, we randomized transplanted mice to receive either 8 mg/kg of anti-PD-1 monoclonal antibody or isotype control intraperitoneally three times a week for two consecutive weeks. Anti-PD-1 treatment dramatically reduced tumor growth and greatly extended the median survival of mice bearing *Dpy30*-KO tumors, with 60% of these animals showing complete tumor regression (**Fig. 6a-c**).

**Figure 6.**
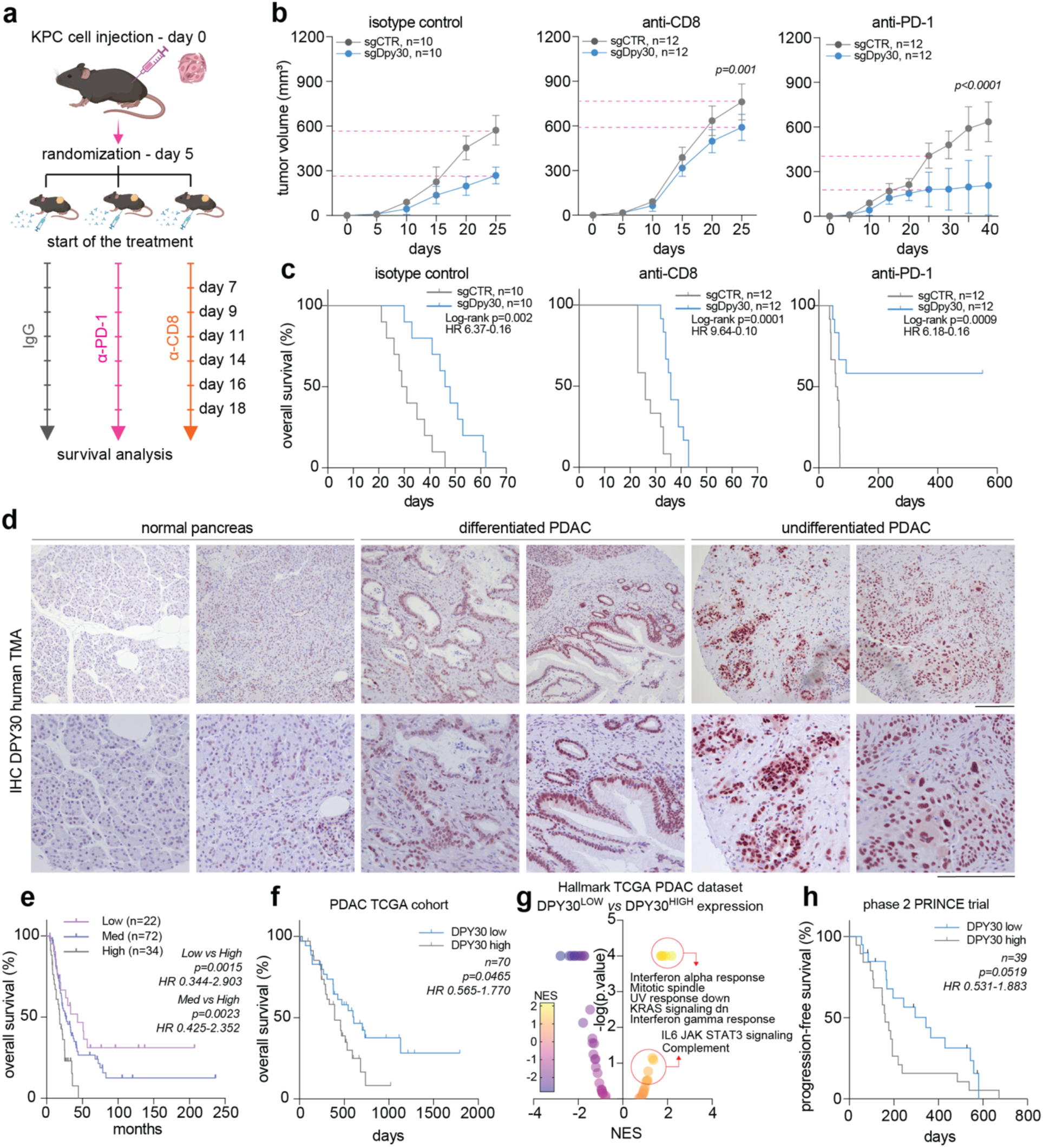
DPY30 expression correlates with tumor stage, poor outcome, and resistance to ICB. **a.** Schematic representation of the in vivo experiment: control (sgCTR) or *Dpy30*-KO (sgDpy30) KPC cells were injected subcutaneously into C57BL/6 mice. After 5 days mice were randomized and treated starting from day 7 with monoclonal antibodies against CD8 (α-CD8) or PD-1 (α-PD-1) or isotype control three times per week, for two weeks. **b-c.** Graphs reporting the tumor volumes (b) or Kaplan-Meier survival curves (c) of sgCTR or sgDpy30 KPC tumors as described in a. **d.** DPY30 immunohistochemical (IHC) staining in normal pancreas and PDAC samples included in the tissue microarray (TMA). **e.** Kaplan−Meier survival curve of patients with PDAC included in the TMA described in (d) and divided according to low (IHC score 0 or 1), medium (IHC score 2), or high (IHC score 3) DPY30 expression. **f.** Kaplan−Meier survival curve of patients in the TCGA PDAC cohort and divided according to low or high *DPY30* expression. **g.** Volcano plot representing GSEA in high-purity TCGA PDAC samples and divided according to high--or low *DPY30* expression. **h.** Kaplan−Meier progression-free survival curve of patients included in the phase 2 PRINCE PDAC trial and divided according to low or high *DPY30* expression. In (c, e, f, h) statistical significance and Hazard Ratio (HR) were calculated using the Log-rank test. In (b) data are presented as mean ± SD and statistical significance was calculated using one-way ANOVA.

To explore the translational relevance of our findings, we next evaluated *DPY30* expression in PDACs by performing immunohistochemical (IHC) analysis on a tissue microarray (TMA) comprising of human PDAC (*n* = 128) and matched tumor-adjacent normal pancreas tissue samples (*n* = 128) collected at the University of Texas MD Anderson Cancer Center. We found that *DPY30* expression was low in normal pancreatic tissues but high in PDAC, particularly in poorly differentiated tumors (**Fig. 6d**). Moreover, we showed that low *DPY30* expression was associated with better survival in the cohort of patients included in the TMA (**Fig. 6e**). To further investigate this association, we interrogated The Cancer Genome Atlas (TCGA) PDAC database and found that, compared to other WRAD core members, high *DPY30* expression was significantly associated with shorter overall patient survival (**Fig. 6f, Extended Data Fig. 10a-c**). Based on these clinical findings, we hypothesized that DPY30 expression in PDAC may be modulated over tumor progression. To test this, we collected pancreas tissues from the KPC mouse model at different time points (46, 68, 69). In support of our hypothesis, IHC analysis showed low DPY30 expression in normal pancreas and metaplastic lesions but a striking increase in expression in transformed cells that became increasingly higher as the lesions progressed from pancreatic intraepithelial neoplasia (PanIN) to high-grade PDAC (Extended Data Fig. 10d). We also noticed that, in normal pancreas, DPY30 was mostly expressed in the endocrine compartment, in line with its role in endocrine cell functional maturation (38, 39), while being weakly detectable in pancreatic ductal and acinar cells (Extended Data Fig. 10d). These data provide evidence that DPY30 expression is associated with PDAC progression in both human and mouse.

To examine whether DPY30 may similarly affect genome integrity and the immune response in human tumors, we queried the transcriptomic data of the TCGA PDAC patient cohort. GSEA analysis of 70 samples with at least 50% purity distributed according to high or low *DPY30* expression showed that low *DPY30* expression was associated with increased expression of genes involved in the complement response as well as in interferon and IL6/STAT3 signaling pathways (**Fig. 6g**). We corroborated our data by querying the transcriptomic data of 47 PDAC patient-derived xenografts (PDXs) established at UT MD Anderson Cancer Center PDACs. GSEA analysis showed that PDAC PDXs expressing low levels of *DPY30* are enriched in genes belonging to the interferon alpha and gamma response pathways (Extended Data Fig. 10e). Finally, we interrogated the dataset of the phase 2 PRINCE trial, in which the efficacy of the PD-1 inhibition (nivolumab) was assessed in patients with metastatic PDAC (70). Strikingly, PDAC patients expressing low DPY30 levels showed a prolonged progression-free and overall survival compared to PDAC patients with high DPY30 expression (**Fig. 6h, Extended Data** Fig. 10f-g). Taken together, our data strongly suggest that DPY30 has a role in regulating DNA replication in PDAC cells, and that low levels of DPY30 favor aberrant DNA replication events that activate an interferon-dependent response that recruits effectors of the adaptive immune response.

## Discussion

While many studies have focused on the genetics and transcriptional fingerprint of PDAC to designate subtype classification schemes for this highly heterogeneous disease, these molecular subtypes have not yet consistently translated to clinical implications (28, 71–73). Cumulative evidence sheds light on the intricate mechanisms underpinning PDAC tumorigenesis and disease progression, which involve context- and time-dependent interactions between genomic and epigenomic scars (74, 75). Recently, a study of the epigenome of PDAC PDXs demonstrated that epigenetic fingerprints are associated with discrete transcriptional signatures and clinical features, suggesting a convergence between genetics, epigenetics, tumor-intrinsic, and tumor-extrinsic features (76).

The disruption of evolutionarily conserved epigenetic complexes, including COMPASS, has been implicated in developmental disorders and cancer, as strict regulation of the activity of these complexes are critical for developmental and physiological processes (37, 48). For instance, in PDAC, the cumulative incidence of mutations in epigenetic regulators is approximately 35%, including recurrent mutations in *KMT2B*, *KMT2C*, *KMT2D*, *SETD1A*, and *SETD1B* (26–30). Interestingly, *DPY30* mutations have not been observed in any cancer.

While the WRAD core of COMPASS complexes is known to regulate the maintenance of epigenetic information in normal and cancer cells, numerous studies have demonstrated non-canonical functions of the WRAD subunits that include transcriptional regulation-independent functions that regulate cell cycle progression (33–35, 49, 77–81). Here, we demonstrate a novel function for the WRAD core as an effector in DNA replication and fork stability, and we show that this function is independent of the core’s function as an MLL/Set methyltransferase (38–40, 42–44). Although loss of DPY30 did not affect the proliferative ability of PDAC cells compared to the inactivation of other WRAD core members, we show that DPY30 integrity is required to prevent excessive DNA replication and maintain genome integrity. Specifically, molecular analyses revealed that DPY30 loss leads to uncoordinated DNA replication, resulting in increased replication stress and, ultimately, promoting genome instability. Moreover, *DPY30*-depleted cells displayed a rapid and unchecked transition from S phase to G2/M and were observed to progress through the cell cycle with mis-segregated chromosomes and persistent DNA damage.

The S-to-G2 cell cycle checkpoint is enforced by the ATR kinase, which governs the coupling of DNA replication with mitotic progression by repressing the activity of CDK1 and ensuring proper timing of G2-specific events (82–84). DNA re-replication can itself activate ATR, which in turn arrests DNA replication and recruits DNA repair effectors (13, 15, 25). However, the biological mechanisms by which this ATR-mediated checkpoint can sense the completion of DNA replication and distinguish this event from DNA over-replication remain unknown. Our findings shed light on how tumor cells with oncogenic signaling and deficient cell cycle checkpoints employ independent, novel pathways to restrain DNA replication to evade cell death mechanisms. Clarifying the interplay between DNA re-replication control, checkpoint activation, and repair of re-replication-associated DNA lesions is fundamental for expanding our knowledge of the biology driving cancer cells.

Whole-genome duplication and CIN frequently occur during the progression of PDAC, and especially at late and metastatic stages. In particular, chromothripsis, a catastrophic mitotic event driving multiple concurrent chromosome structural alterations, is frequently observed in advanced PDAC (30). Interestingly, *DPY30* expression is upregulated during tumorigenesis, and poorly differentiated PDACs displayed the highest *DPY30* expression. This suggests that DPY30 may somehow buffer PDAC cells from oncogene-driven uncontrolled replication and the detrimental effects of CIN. All models evaluated in our study were *P53* deficient, thus warranting additional investigations to determine whether *P53* inactivation represents a specific genetic context in which DPY30 function is essential for regulating DNA homeostasis. Further, because aneuploid cells are characterized by deregulated metabolic programs and protein networks, future studies should assess whether DPY30 loss, through induction of aneuploidy, can evoke cellular adaptive responses, which may ultimately be exploited as co-lethality and druggable targets (1, 6, 85).

Overall, our study demonstrates that DPY30 loss leads to the accumulation of PDAC cells with complex karyotypes and increased TMB, which, in turn, results in the selective clearance of damaged proliferating cells by CD8+ T cells. Considering the paucity of treatment options for patients with PDAC, our work provides evidence that PDAC with low *DPY30* expression may respond to combined therapy of immune-checkpoint inhibitors and conventional chemotherapy.

## Methods

### Patient-derived samples

Patient-derived samples were obtained from consented patients under an Institutional Review Board (IRB)-approved protocol LAB00-0396 chaired by Dr. Michael Kim. (UTMDACC).

### Cell Biology Experiments

#### Cell culture

Human 293T and RPE-1 (ATCC), PDX-derived PATC53 (PDX1), PATC66 (PDX2), and mouse-derived PDAC cells KP^-/-^C (internally registered as WY5549 and derived from *Ptf1aCre;LSL-Kras^G12D/+^;LSL-Trp53^L/L^*mouse model) and KPC (internally registered as N150F and derived from *Ptf1aCre;LSL-Kras^G12D/+^;LSL-Trp53^R172H/R172H^* mouse model, and named as N150F, see below “mouse strains were cultured in Dulbecco modified Eagle medium (DMEM, Sigma) supplemented with 10% fetal bovine serum (FBS, Gibco), 100IU/mL Penicillin (Gibco) and 100μg/mL Streptomycin (Gibco).

#### Lentiviral production

Viral particles were produced using packaging plasmids psPAX2 and pMD2.G. Briefly, 293T cells (ATCC) were seeded in 15cm diameter plate in DMEM supplemented with 10% FBS (Gibco), 100IU/mL Penicillin (Gibco), 100μg/mL Streptomycin (Gibco), 2 mM Glutamine (Gibco) and allowed to adhere. Transfection mix was prepared by adding the vectors of interest in Opti-MEM (Gibco) in presence of Polyethyleneimine (PEI). Supernatant was collected 72 hours after transfection, filtered through 0.45μm low-protein binding filters (Corning) and concentrated in sterile PBS (Sigma Aldrich) after ultracentrifugation at 25,000 rpm for 2 hours. Concentrated virus was freshly used to transduce target cells.

#### Cell transduction with lentiviral particles

Human and mouse cells were transduced overnight with concentrated lentiviral particles (see Virus preparation) in the presence of 8 μg/mL polybrene. After 72 hrs from infection, the culture medium was replaced with a fresh medium supplemented with antibiotic selection (G418 50 μg/mL or puromycin 1-5 μg/mL). Overexpression, down-regulation, or knockout of the targets were evaluated by western blot after one week of antibiotic selection. For doxycycline-inducible shRNAs, after one week of antibiotic selection, cells were pulsed with doxycycline hyclate (Millipore-Sigma, D5207) 1 μg/mL at different time points, as indicated in figures and gene expression was evaluated by western blot.

#### Cell transfection

Cells were seeded at 50% confluency in 10 cm diameter plate in complete medium and allow to adhere overnight. Transfection mix was prepared by adding the 10 μg of the indicated plasmids in 600 μL Opti-MEM (Gibco) in presence of 40 μL polyethyleneimine (PEI). Cells were then transfected for 8-12 hrs and protein expression was evaluated using Western Blot.

#### Generation of barcoded KPC (N150F) cell line

KPC cells were infected overnight with CloneTracker™ XP Lentiviral Expressed Barcode Library (Cellecta) in presence of polybrene (8μg/ml) (MOI < 0.1). After infection and puromycin selection, cells were passaged for 10 generations before injection into NSG or C57BL/6 mice.

#### Cell proliferation and clonogenic assay

Human and mouse cells were seeded in a 96-well plate (500-1000 cells per well) in medium supplemented with doxycycline 1 μg/mL or vehicle. After cell adhesion, cells were incubated at at 37°C 5%CO_2_ in the IncuCyte (Essenbioscience) incubator. Cell confluency was measured and analyzed over a period of 5-8 days using and medium was changed every 48 hrs.

For clonogenic assay, 100-500 cells were seeded in a 6-well plate in medium supplemented with doxycycline 1 μg/mL or vehicle and maintained at 37°C 5%CO_2_ (medium was changed every 48 hrs). After 8-15 days, clones were fixed and stained with crystal violet (0.25% crystal violet in methanol 20%). Colonies with more than approximately 50 cells were counted manually and clonogenic survival fraction was expressed as the relative plating efficiencies of the irradiated cells to the control cells.

### Molecular Biology Experiments

#### Short-hairpin RNAs (shRNAs) cloning

For gene silencing, shRNAs from Millipore-Sigma were cloned into the constitutive pLKO.1-TRC cloning vector according to the manufacturer’s protocol. These same shRNA sequences were cloned into pENTR-THT and integrated into the inducible Tet-Puro backbones.

#### Single-guide RNAs (sgRNAs) design and cloning

sgRNAs were designed with “GenScript CRISPR sgRNA Design Tool” (https://www.genscript.com/gRNA-design-tool.html?a=post). First, 5’-phosphorilated oligos were annealed and diluted 1:20. Then 1 μL of each annealed and diluted sgRNA was cloned in digested lentiCRISPR-V2 according to Dr. Feng Zhang’s protocol (https://media.addgene.org/cms/files/Zhang_lab_LentiCRISPR_library_protocol.p df). NEB® Stable Competent E. coli (C3040I) colonies resistant to ampicillin antibiotic selection were amplified, and presence of sgRNA was confirmed by Sanger sequencing. Positive clones were transfected individually in 293T cells along with vectors for lentiviral packaging production. Human or mouse cells were infected by lentiviral particles carrying a specific sgRNA and selected for puromycin resistance. Cut efficiency of sgRNA was tested by T7 Endonuclease I (NEB #M0302L) assay on the DNA of infected cells, according to the manufacturer’s protocol (https://www.neb.com/protocols/2014/08/11/determining-genome-targeting-efficiency-using-t7-endonuclease-i). The sgRNA sequences for knocking out DPY30 are the following: GAGCCAGAGCAGATGCTGGA (sgDPY30#1), GGGACTTGCTGTGCTTGCAA (sgDPY30#2) for human and GAGTCGGAGCAGATGCTGGA (sgDpy30) for mouse.

#### Plasmids

The following vectors were purchased from Addgene: psPAX2 (#12260), pMG2.D (#12259), lentiCRISPR-V2 (#52961), pLKO.1-TRC (#8453), pENTR-THT (#55790), Tet-Puro (#21915), pLenti-PGK-Neo-PIP-FUCCI (#118616), emEGFP-Mcm2 (mouse, #54163), emEGFP-Mcm3 (mouse, #54165), emEGFP-Mcm4 (mouse, #54167), emEGFP-Mcm5 (mouse, #54169), emEGFP-Mcm6 (mouse, #54171), emEGFP-Mcm7 (mouse, #54173), pcDNA3 Flag-hDPY30 (#15554), pcDNA3 Flag WDR5 (#15552), pCMV-8xHis-Ub (#107392). The following vectors were purchased from Genscript: pcDNA3.1 WDR5_L240K(+)-N-HA (L240K WDR5_OHu17370C), pcDNA3.1 WDR5_V268E(+)-N-HA (V268E WDR5_OHu17370C), pcDNA3.1 WDR5-(+)-N-HA (WDR5_OHu17370C), pcDNA3.1(+)-C-HA-L69D-hDPY30 (L69D DPY30_OHu17211C), pcDNA3.1(+)-C-HA-hDPY30 (DPY30_OHu17211C), pcDNA3.1(+)-C-HA-mdpy30 (Dpy30_OMu07274C), pcDNA3.1(+)-N-6His-hMCM2 (MCM2_OHu09867C), pcDNA3.1(+)-N-6His-hMCM3 (MCM3_OHu107378C), pcDNA3.1(+)-N-6His-hMCM4 (MCM4_OHu11591C), pcDNA3.1(+)-N-6His-hMCM5 (MCM5_OHu02170C), pcDNA3.1(+)-N-6His-hMCM6 (MCM6_OHu20108C), pcDNA3.1(+)-N-6His-hMCM7 (MCM7_OHu28012C). The following vectors were purchased from Sigma-Millipore: shCTR (SHC002), shDPY30#1 (TRCN0000297810), shDPY30#2 (TRCN0000280267), shWDR5#1 (TRCN0000118047), shWDR5#2 (TRCN0000118049), shRBBP5#1 (TRCN0000165777), shRBBP5#2 (TRCN0000165100), shASH2L#1 (TRCN0000019274), shASH2L#2 (TRCN0000019276), shDpy30#1 (TRCN0000034420), shDpy30#2 (TRCN0000034419).

#### Protein extraction, Immunoprecipitation and Western Blot Analyses

For cellular protein lysates, cells were scraped on ice using cold Ripa lysis buffer (150 nM NaCl, 50 mM Tris HCl pH 8, 1% Igepal, 0.5% sodium deoxycholate, 0.1% SDS - Boston BioProducts, BP-115) supplemented with a HALT protease and phosphatase inhibitor cocktail (ThermoFisher). Cell lysates centrifuged at 13,200 rpm for 20 min at 4°C and supernatants were collected.

For immunoprecipitates (IPs), cells were scraped using NP40 buffer (10 mM NaCl, 10 mM Tris HCl pH 7.5, 0.5% Nonidet P-40 - Boston BioProducts, BP-119) supplemented with a HALT protease and phosphatase inhibitor cocktail (#78440, ThermoFisher Scientific). Cell lysates were sonicated twice at 30% amplitude for 15 sec on ice and centrifuged at 13,200 rpm for 20 min at 4°C and supernatants were collected. Immunoprecipitation was performed by binding the primary antibody to protein G magnetic beads (Dynabeads, ThermoFisher Scientific, 10004D) for 30-60 min at RT. Bs3 (A39266, ThermoFisher Scientific) was used to crosslink primary antibodies to protein G. Cell lysates were diluted with PBS-T buffer (PBS-0.1% Tween20) supplemented with protease/phosphatase inhibitor, added to antibody-bead complexes and incubated overnight at 4°C. IPs were then washed in PBS-T buffer, resuspended in 4X NuPage Sample Buffer (Invitrogen) supplemented with 1% β-mercaptoethanol.

IPs and proteins were separated in 4-20% SDS-PAGE (Criterion Precast Midi Gel, Bio-Rad) and transferred to nitrocellulose membranes (Trans-Blot Turbo Midi 0.2 μm nitrocellulose transfer pack, Bio-Rad). Membranes were blocked with 5% non-fat dried milk in PBS-T and incubated at 4°C overnight with primary antibodies.

Membranes were washed in PBS-T and incubated 1 hour at RT with the appropriate horseradish peroxidase-conjugated secondary antibodies (Cell Signaling Technology) for ECL detection (SuperSignal™ WEST Pico PLUS Chemiluminescent Substrate, ThermoFisher) or with the appropriate IRDye 800CW or IRDye 680RD Dye-Labeled secondary antibodies (LICORbio). Band quantification was performed using the QuantiONE software (Bio-Rad Laboratories). The Re-Blot Plus Strong Solution (#2404, EMD Millipore) was used to strip the membranes, when reblotting was needed.

#### Purification of proteins on newly synthesized DNA (iPOND)

iPOND was performed essentially as described previously (86). Briefly, 293T or KPC cells were seeded in a square at 70% confluency (one square per each iPOND sample). Cells were pulsed with 10 μM EdU (5-Ethynyl-2’-deoxyuridine, Millipore-Sigma, #900584), for 5 and 10 min to enable the detection of protein bound to active DNA replication forks. For each experiment, two negative controls were included: one sample was not pulsed with EdU and one sample was pulsed with EdU followed by 60 min thymidine chase. All samples were fixed with 1% formaldehyde (ThermoFisher Scientific) for 20 min at RT, and quenched using 125 mM Glycine (Millipore-Sigma) solution for 5 min at RT. After fixation, samples were collected in 50-ml tube and washed (once with PBS+1% BSA and once with PBS). Samples were permeabilized using PBS+0.1% Tryton-X100 for 10 min in gentle shaking at RT and transferred in a 15-ml tube. Copper(ii) Sulfate (Millipore, Sigma #451657), (+)-sodium L-ascorbate (Millipore, Sigma #A7631) and Biotin Azide (Click chemistry tools, Fisher) were used to perform the Click-reaction for biotin-conjugate the incorporated EdU. After washing steps, cell pellets were resuspended in SDS 1% for protein extraction and sonicated 3 times at 30% amplitude for 30 sec on ice. After centrifugation, cleared lysates were diluted 1:2 with PBS and % 1% of protein lysate was kept as input control. Cleared lysates were then incubated with Pierce™ Streptavidin Magnetic Beads (ThermoFisher Scientific, 88817) for 2 hr at RT in gentle shaking. Protein complexes were washed three time with PBS-T and purified proteins were resuspended in 4X NuPage Sample Buffer (Invitrogen) supplemented with 1% β-mercaptoethanol.

#### In vitro immunofluorescent staining

Cells were plated at 50% of confluency over a round glass coverslip (Chemglass Life Sciences, cat. n. CLS1763012), allowed to adhere overnight, treated with doxycycline as appropriate to induce gene silencing and fixed using 4% paraformaldehyde for 15 min at RT. Cells were then washed twice in PBS and permeabilized (0.1% triton X100 in PBS) for 15 min at RT. After a second round of washes in PBS, cells were blocked using blocking buffer (3% BSA, 3% goat serum, 0.01% triton X100 in PBS) 1h at RT. Primary antibodies were diluted in blocking buffer and incubated overnight at 4°C. After washes, secondary antibodies (conjugated with Alexa-488, Alexa-594, Alexa-647 or Alexa-750) were diluted in blocking buffer and incubated 1h at RT. DAPI was used to counterstain nuclei. Images were captured with a Hamamatsu C11440 digital camera, using a wide-field Nikon Ni-E fluorescent microscope.

#### Proximity Ligation Assay (PLA) and SIRF assay

For PLA, Duolink® In Situ Detection Reagents Red (Millipore-Sigma) were used and cells were stained according to the manufacturer protocol. Briefly, cells were plated, fixed and blocked as described above. After primary antibody incubation, cells were washed with Buffer A and then the Duolink® In Situ PLA® probe anti-rabbit or anti-mouse were added as appropriate. After ligation and amplification reactions, cells were imaged using Zeiss LSM880 Airyscan confocal microscope (63X objective) at the Advanced Microscopy Core (MD Anderson Cancer Center). SIRF assay was performed essentially as described in (52, 53) with minor modifications. Cells were plated on coverslips as described above and were pulsed with 20 μM BrdU for 5 min. The primary antibodies used were anti-WDR5 (rabbit), anti-DPY30 (rabbit, Bethyl), anti-PCNA (rabbit) and anti-BrdU (mouse). Click-IT reaction was not performed.

#### DNA Fiber Assay

Fiber stretching was performed essentially as described in (87). Briefly, cells were double-pulsed precisely for 20 min with 25 μM CldU (Chloro-2’-deoxyuridine (Millipore-Sigma, C6891) followed by 250 μM IdU (5-Iodio-2’-deoxyuridine, Millipore-Sigma, I7125). Cells were harvested and diluted at 2×10^5^cells/ml. A 3-μL drop of cell suspension was allowed to air-dry in a glass slide and fiber spreading was carried out by adding 10 μL spreading buffer (0.5% SDS in Tris-HCl EDTA). Fibers were fixed in Methanol/Acetic Acid 3:1 10 min, allowed to completely dried and rinsed with deionized water. DNA was denatured with HCl 2N for 1.5 hr ar RT. After washing steps, slides were blocked in blocking buffer (PBS 2% BSA, 2% goat serum, 0.1% Tween20) 1 hr at RT. CldU and IdU were recognized by using the following primary antibodies: rat monoclonal anti-BrdU clone BU1/75 and mouse monoclonal anti-BrdU clone B44. The antibodies were diluted in blocking buffer and incubated 4 hr at RT, followed by secondary antibody (goat anti-rat IgG (H+L) Alexa Fluor® 488, goat anti-mouse IgG (H+L) Alexa Fluor® 594, ThermoFisher Scientific). Imaging was performed on the Leica SP8 laser scanning confocal microscope at the Advanced Microscopy Core (MD Anderson Cancer Center).

#### Chromosome mis-segregation

For the *in vitro* analysis, cells were plated at 50% of confluency over a round glass coverslip (Chemglass Life Sciences, cat. n. CLS1763012), allowed to adhere overnight and fixed using 4% paraformaldehyde for 10 min at RT. DNA was stained using DAPI 10 min at RT. Chromosome mis-segregation were evaluated using a wide-field Nikon Ni-E fluorescent microscope (40X objective). For *in vivo* analysis, H&E samples from human or mouse-derived tumors grown in recipient mice were evaluated for anaphases with chromosome mis-segregation events using a wide-field Nikon Ci-L plus light microscope (100X objective). Analyses was performed by evaluating chromosome mis-segregation in H&E samples (5 mice per group, one section per mouse), and for each mouse at least 20 anaphases were counted.

#### Preparation of Metaphase Spreads

Analysis of chromosomal aberrations was performed at the Molecular Cytogenetics Facility (MD Anderson Cancer Center) as previously described (32). Cells were incubated with Colcemid (KaryoMAX™, ThermoFisher Scientific, #15212012) at 0.04 μg/mL for 3 hours, trypsinized, washed, and spun down. Cells were incubated at RT for 15 minutes in hypotonic solution (0.075M KCl). Cells were fixed by adding three times 2 mL of fixative solution (glacial MeOH + acetic acid 3:1), followed by centrifugation. Cell suspension was dropped on glass slide. Slides were allowed to air-dry, stained with 4% Giemsa staining solution (KaryoMAX™, ThermoFisher Scientific, #10092013) for 5 min and mounted. At least 35 metaphases were analyzed from each sample for chromosomal aberrations. Images were captured using a Nikon 80i microscope equipped with karyotyping software from Applied Spectral Imaging, Inc.

#### Cell cycle synchronization and analysis by flowcytometry

Human and mouse cells were seeded in a 6-cm diameter plate (1×10^5^ cells per plate) in complete medium and allowed to adhere to plate for 16 hr. To synchronize cells in early S phase, a double thymidine (Millipore-Sigma, T9250) pulse was used. Briefly, cells were cultured in presence of 2 mM thymidine for 18 hrs, released in complete medium for 8 hrs and arrested in S phase with a second thymidine-pulse for 18 hrs. Cells were then released at different time point as indicated in figure using complete medium. To synchronize cells in G2/M phase, cells were cultured in presence of 2 mM thymidine for 18 hrs, released in complete medium for 8 hrs and arrested in G2/M phase with 100 ng/ml Nocodazole (Millipore-Sigma, M1404) for 16 hrs. For DNA content analyses cells were harvested at specific time point, resuspended in 250 μL of ice-cold PBS and fixed by adding 750 μL of absolute ethanol dropwise while vortexing. Cells were kept an overnight in fixative solution, washed twice with PBS and DNA stained using DAPI. Data were acquired using BD Celesta analyzer and analyzed by FlowJo.

#### BrdU incorporation assay and Re-replication analyses using flow cytometry

Human and mouse cells were seeded in a 6-cm diameter plate (1×10^5^ cells per plate) in complete medium and allowed to adhere to the plate for 16 hr. Cells were pulsed with 15 μM BrdU for 2 hrs, harvested and fixed using APC or FITC BrdU kit (#552598 or #559619 BD bioscienses) following manufacturer protocol. Alternatively, human PDAC cells were fixed with ethanol as described above and DNA was denaturated using 2N HCl solution for 20 min. Cells were then washed three time with PBS, blocked in PBS 5% BSA and PE-conjugated anti-BrdU antibody was incubated 1 hr at RT. After washing with PBS, nuclei were counterstained using DAPI. Data were acquired using BD Celesta analyzer and analyzed by FlowJo.

#### Cell sorting

Human cells were seeded in a 15-cm diameter plate at 70% confluency in complete medium and allowed to adhere to the plate for 24 hr. Cells were harvested and resuspended in FACS buffer (3% FBS, 1mM EDTA) and DAPI was used as live/dead reagent. Cells were sorted using Influx Silver sorter and data were analyzed by FlowJo.

#### Immunophenotyping using flow cytometry

After tumor dissociation and single cells extraction (see Cell isolation from tumors), staining was performed essentially as described in (88). Briefly, cell surface antigens were stained with the appropriate primary antibodies for 20 minutes at 4°. For the staining of intracellular proteins, samples were washed, fixed, permeabilized for 30 min at RT and subsequently incubated with the appropriate primary antibodies for 1 hr at RT.

Dead cells were stained using the fixable viability violet dye Zombie Red (Invitrogen) for 10 min at RT followed by the blocking of Fc receptors with TruStain fcX (Biolegend) for 15 min at 4°C. Cells were analyzed on LSRFortessa X-20 flow cytometers (BD Biosciences) and data were analyzed with FlowJo software version 10.8.2.

#### Ex vivo stimulated cytokine secretion

Single-cell suspensions from tumors were resuspended in RPMI 1640 with 10% FCS, were added to anti-CD3 (clone 145-2C11)/anti-CD28 (clone 37.51) antibody-coated (overnight at 10 μg/ml antibody) culture plates for 4hr at 37°C in the presence of 1 μg/ml Golgiplug (BD Bioscience) and Monensin (Biolegend) and processed for TNF and IFNγ intracellular cytokine staining.

#### Histopathology, immunohistochemistry, and immunofluorescence

Tumor specimens were fixed overnight in 4% buffered paraformaldehyde, transferred to 70% ethanol, and then embedded in paraffin using Leica ASP300S processor. For histopathological analysis, tumors were sectioned (Leica RM2235, 5μm thickness) and serial slides were collected. For every series one section was stained with hematoxylin and eosin and remaining sections were kept for either immunofluorescence or immunohistochemical analysis. For Alcian Blue (Vector Laboratories, H3501) and Masson’s Trichrome staining (VitroView™ Masson’s Trichrome Stain Kit, VB-3016), samples were processed according to the manufacturers’ protocols.

Histological samples were processed as previously described (74). Briefly, after cutting, baking, and deparaffinization, sections underwent antigen retrieval using Citra-Plus Solution (BioGenex) according to specifications.

*For immunohistochemistry (IHC) staining*, endogenous peroxidases were inactivated by 3% hydrogen peroxide and non-specific signals were blocked using 5% BSA, 5% goat serum, and 0.1% Triton X-100. Primary antibodies were applied and incubated overnight at 4°C. ImmPRESS HRP IgGs were used as secondary antibodies and ImmPACT NovaRED® (Vector Laboratories, # SK-4805) was used for detection. Images were captured with a Nikon DS-Fi1 digital camera using a wide-field Nikon Ci-L plus light microscope.

Tissue Microarray slides of PDAC samples with available survival data (n=128) collected at MD Anderson Cancer Center were stained for DPY30 as described above. DPY30 nuclear expression was evaluate and categorized in blind as follow: Low (score 0=negative or score 1=weak), medium (score 2=moderate) and high (score 3=strong) expression. Kaplan-Meier survival curve was generated by clustering PDAC patients according to DPY30 expression and log-rank test was used to compare the survival difference among different groups.

*For immunofluorescence staining*, secondary antibodies conjugated with Alexa-488, Alexa-594, Alexa-647 or Alexa-750 were used. DAPI nuclear staining (#40043, Biotium) was used to counterstain nuclei. Images were captured with a Hamamatsu C11440 digital camera, using a wide-field Nikon Ni-E fluorescent microscope or Vectra Polaris (Akoya Biosciences).

#### Cyclic immunofluorescence

sAfter the first staining and image acquisition using PE Vectra polaris, coverslips were removed by incubating slides with 37°C PBS for 20 min. Slides were then incubated with stripping buffer (62.5 mM Tris Base / Tris-HCl, 2% Sodium Dodecyl Sulfate, 0.8% Beta ME, 55°C, 30 min), and then stained with another round of antibodies. Slides were washed with PBS 0.1% Triton-×100 between each step. DAPI nuclear staining (#40043, Biotium) was used every cycle.

#### Antibodies

Primary antibodies purchased from Cell Signaling Technology: anti-WDR5 (clone D9E1I, #13105), anti-Cleaved Caspase-3 (Asp175, #9661), anti-STING (clone D2P2F, #13647), anti-STING (S365) (clone D1C4T, #62912), anti-ASH2L (clone D93F6, #5019), anti-RbBP5 (#8037), anti-MCM2 (S139) (clone D1Z8X, #12958), anti-MCM2 (clone D7G11, #3619), anti-MCM3 (#4012), anti-MCM4 (clone D3H6N, #12973), Rabbit monoclonal anti-MCM7 (clone D10A11, #3735), anti-Histone H3 (pS10) (#9701), anti-Cdc6 (clone C42F7, #3387), anti-H3K4Me3 (clone C42D8, #9751), anti-His Tag (clone 27E8, #2366). Primary antibodies purchased from Millipore-Sigma: anti-DPY30 (#HPA043761), anti-γH2AX (pS139) (clone JBW301, #05-636), anti-beta-Actin (clone AC-74, #A5316). Primary antibodies purchased from Bethyl Laboratories: anti-DPY30 (#A304-296A), anti-MCM5 (#A300-195A), anti-MCM6 (#A300-194A), anti-WDHD1 (#A301-140A), anti-RPA2 (pS4-8) (#A300-245A). Primary antibodies purchased from Santa Cruz Biotechnology: anti-WDR5 (clone G-9, #sc-393080), anti-Cyclin E (clone HE12, #sc-247), anti-Histone H3 (clone 1G1, #sc-517576), anti-Geminin (clone FL-209, #sc-13015), anti-Luciferase (clone C-12, #sc-74548). Primary antibodies purchased from Abcam: anti-BrdU (clone BU1/75 – ICR1, #ab6326), anti-H3K4Me1 (#ab8895), anti-H3K4Me3 (#ab8580), anti-H3K4Ac (#ab4729), Alexa Fluor® 594 anti-αSMA (clone 1A4, #ab202368), Alexa Fluor® 488 anti-αSMA (clone 1A4, #ab184675). Primary antibodies purchased from ThermoFisher Scientific: anti-PCNA (#PA5-16797), anti-Cyclin A2 (clone E23.1, #MA1-154), anti-GFP (clone GF28R, #MA5-15256), anti-Ki67 (#MA5-14520), anti-HA tag (clone 2-2.2.14, #26183), anti-CD8a (clone 4SM15, #14-0808-82), PE-conjugated anti-BrdU (clone BU20A, #12-5071-42). Other primary antibodies: anti-BrdU (clone B44, #347580, BD Laboratories), anti-Cdc6 (pS54) (#102-15286, RayBiotech), anti-Ki67 (#14965-1-AP, Proteintech), anti-Flag tag (clone OTI4C5, #TA50011-100, OriGene Technologies), anti-CD3 (#A045201-2, Agilent Technology), Rabbit gamma globulin (#31887, Invitrogen). Antibodies purchased from Biolegend: anti-mouse Ki67 (clone 16A8, #652402), BV 510 anti-mouse CD8a (clone 53-6.7, #100751), PE anti-CD279 (PD-1) (clone RMP1-30, #109104), PB anti-Granzyme B (clone GB11, #515408), PE-Cy5 anti-CD3 (clone 17A2, #100274), Pe-Cy-7 anti-Foxp3 (clone FJK-16S, #25-5773-82), APC-Cy7 anti-CD4 (clone GK1.5, #100414), Alexa Fluor® 700 anti-CD45 (clone 30-F11, #103128), PerCP-Cy5.5 anti-Thy1.2 30-H12 (clone 30.H12, #105338), BV785 anti-NK1.1 (clone PK136, #108749), Alexa Fluor® 647 anti-Tnf (clone MP6-XT22, #506314), Alexa Fluor® 700 anti-Infγ (clone XMGG1.2, #505824). Secondary antibodies purchased from Vector Laboratories: ImmPRESS® HRP goat anti-rabbit IgG (#MP-7451), ImmPRESS® HRP goat anti-mouse IgG (#MP-7451). Secondary antibodies purchased from ThermoFisher Scientific: Goat anti-rabbit IgG (H+L) Alexa Fluor® 488 (#A-11008), Goat anti-rabbit IgG (H+L) Alexa Fluor® 594 (#A-11012), Goat anti-rabbit IgG (H+L) Alexa Fluor® 647 (#A-21244), Goat anti-rabbit IgG (H+L) Alexa Fluor® 750 (#A-21039), Goat anti-mouse IgG (H+L) Alexa Fluor® 488 (#A-11001), Goat anti-mouse IgG (H+L) Alexa Fluor® 594 (#A-11005), Goat anti-mouse IgG (H+L) Alexa Fluor® 647(#A-21235), Goat anti-rat IgG (H+L) Alexa Fluor® 488 (#A-11006), Goat anti-rat IgG (H+L) Alexa Fluor® 647 (#-21247).

#### RNA extraction

RNA extraction was achieved using Qiagen RNeasy Micro kit following manufacturer’s instruction. Briefly, RNA extraction was performed from KPC and KP^-/-^C cells or tumors stably knocked out for Dpy30 or control and from PATC53 stably expressing doxycycline-inducible scramble control (shCTR) or shRNAs targeting WDR5 or DPY30 pulsed for 48 hrs with doxycycline. Cells grown *in vitro* were scraped using 350 μL of RLT buffer supplemented with β-mercaptoethanol. For RNA extraction from tumors, tissue specimens (approximately 2×2mm) were snap-frozen and kept at −80°C until further processing. Tumors were then weighted and dissociated in RLT buffer supplemented with β-mercaptoethanol using gentleMAX™ M tubes (Miltenyi Biotec) and RNA_02 program. DNA digestion was performed by adding recombinant DNAse I (Millipore-Sigma, #4536282001). to purification column. Total RNA was eluted in RNAse-free water, quantified, and submitted for sequencing.

#### Genomic DNA extraction

DNA extraction was achieved using Qiagen DNaeasy Blood and Tissue kit following manufacturer’s instruction. Briefly, cells grown *in vitro* were harvested and cell pellets were snap-frozen and kept at −80°C. For DNA extraction from tumors, tissue specimens (approximately 2×2mm) were weighted and dissociated by mechanical digestion in 180 μL Buffer ATL supplemented with 20 μL proteinase K stock solution. Total DNA was eluted in DNAse-free water, quantified, and submitted for sequencing.

#### Chromatin immunoprecipitation (ChIP)

ChIP was performed essentially as described in (89) with optimized DNA shearing condition and few modifications. Briefly, 9×10^6^ PATC53 cells stably expressing doxycycline-inducible shRNAs against DPY30, WDR5 or scramble control were seeded in a 15-cm diameter plate in complete medium in the presence or not of 1 μg/ml doxycycline for 48 hrs. Cells were then harvested, resuspended in 3 mL of complete medium, cross-linked with 200μL of 16% formaldehyde for 10 min at 37°C in gentle stirring and quenched with 200 μL Glycine 2M for 5 min at 37°C in gentle stirring. Tissue lysis, sonication and chromatin immunoprecipitation were performed as described in (31, 74).

#### ATAC-seq

Cells were processed as described in (74). Briefly, 100,000 human PATC53 cells were lysed in lysis buffer for 5 minutes on ice. Tn5 tagmentation was carried out for 30 minutes at 37 °C. After indexing and PCR amplification, DNA libraries were multiplexed and sequenced on HiSeq X with 150nt PE configuration.

#### Immuno-precipitate mass spectrometry

KPC, PATC53 and PATC66 cells were seeded in 15cm diameter plate at 50% confluency in complete medium. Cells were scraped in RIPA buffer as described above after 24 hr from cell seeding. For each sample, 2 mg of lysate were immunoprecipitated for DPY30 or control IgG and immunocomplexes were resuspend in 100μL of NuPage sample buffer, and 20μL were resolved using SDS-PAGE for quality control. For proteomics, 40μL of each sample were processed by SDS-PAGE using 10% Bis-Tris NuPage Mini-gel (Invitrogen) with the MES buffer system. The gel was run 1cm and the mobility region excised into 10 equally sized bands. Each band was processed by in-gel digestion with trypsin using a robot (ProGest, DigiLab) with the following protocol: i) washed with 25mM ammonium bicarbonate followed by acetonitrile, ii) reduced with 10mM dithiothreitol at 60°C followed by alkylation with 50mM iodoacetamide at RT, iii) digested with sequencing grade trypsin (Promega) at 37°C for 4h, iv) quenched with formic acid and the supernatant was analyzed directly without further processing. For mass spectrometry, half of each digested sample was analyzed by nano LC-MS/MS with a Waters M-Class HPLC system interfaced to a ThermoFisher Fusion Lumos mass spectrometer. Peptides were loaded on a trapping column and eluted over a 75μm analytical column at 350nL/min; both columns were packed with Luna C18 resin (Phenomenex). The mass spectrometer was operated in data-dependent mode, with the Orbitrap operating at 60,000 FWHM and 15,000 FWHM for MS and MS/MS respectively. The instrument was run with a 3s cycle for MS and MS/MS. 5hrs of instrument time was used for the analysis of each sample.

Quantitative real time PCR. RNA from cells was extracted using RNeasy Mini kit (Qiagen, #74104). Total RNA was quantified using the Nanodrop Spectrophotometer (ThermoFisher). The synthesis of cDNA was performed using the SuperScript VILO Master Mix (ThermoFisher) in an PCR Thermocycler (Bio-Rad). cDNAs were amplified using PowerUp™ SYBR™ Green Master Mix (Invitrogen, A25742) and Real-time PCR Detection System (Applied Biosystems). Normalization of the data was performed using housekeeping genes (*ACTB, GAPDH, 18S*). The following primers (Sigma) were used (5’ – 3’): RFC3 Forward TTCGTGGAAGGCTGTATGAGC, RFC3 Reverse CCTCCCCTTTCAGTTGTCCAT; *GINS2* Forward CAATGCCCAGCCCTTACTACA, *GINS2* Reverse GCCTTCGGGATGTTGTCTGA; *FEN1* Forward GTTCCTGATTGCTGTTCGCC, *FEN1* Reverse ATGCGAATGGTGCGGTAGAA; *IFNL1* Forward GGAGGCATCTGTCACCTTCAA, *IFNL1* Reverse GGACTCAGGGTGGGTTGAC; *IRF-AS1* Forward CACCGCGTCTGTGTCTGTA, *IRF-AS1* Reverse GCCACCTACGATATACCACCC; *TRIM10* Forward TGGCTAACGTGGTGGAGAAC, *TRIM10 Reverse TGCTCTTGGCAGACATCCTC; ACTB* Forward CCAGAGGCGTACAGGGATAG, *ACTB* Reverse CCAACCGCGAGAAGATGA; *18S* Forward CTCAACACGGGAAACCTCAC, *18S* Reverse CGCTCCACCAACTAAGAACG; *GAPDH* Forward GAAGGTGAAGGTCGGAGTC, *GAPDH* Reverse GAAGATGGTGATGGGATTTC.

#### NGS library production for barcode detection in bulk tumors

Frozen tumors were minced and suspended in Buffer P1 (Qiagen, 1 mL buffer per 100 mg tumor) and a scale for cell number quantification was added. This mixture was homogenized in a gentleMACS homogenzier (Miltenyi Biotec). After homogenization, the samples were transferred into 15 mL polypropylene tubes (Falcon) and RNase A (100 μg/mL) was added (10 μL per 100 mg tumor). After 10 minutes, 10% SDS (Promega) and Proteinase K (Qiagen) were added (both 50 μL per 100 mg tumor) and incubated at 56°C for 20 minutes. The DNA was then sheared by passing the lysate through a 23G syringe needle 10 times. The lysates were transferred into 1 mL polypropylene tubes and DNA was purified using Phenol:Chloroform:Isoamyl alcohol (25:14:1 pH 8.0, Sigma Aldrich) and Chloroform:Isoamyl alcohol (24:1, Sigma Aldrich). The aqueous DNA was precipitated by adding 3M sodium acetate (Sigma Aldrich, 90uL per 1 mL sample) and isopropanol (Fisher Scientific, 720 μL per 1 mL sample). DNA was centrifuged at 14,000 rpm for 20 minutes at 4°C and washed with 70% ethanol (Fisher Scientific). Once dried, the DNA pellets were dissolved in Ultrapure distilled water (Thermo Fisher Scientific) and DNA concentration was quantified using NanoDrop 2000 (Thermo Fisher Scientific).

Barcodes were amplified through 2 rounds of PCR on the DNA samples using Titanium Taq DNA polymerase (Clontech, Takara). The first PCR was performed using the following protocol: 3 minutes at 94°C followed by 16 cycles of 30 seconds 94°C, 10 seconds 60°C, and 20 seconds 68°C, followed by final extension at 72°C for 2 minutes then hold at 4°C. The sequences of the primers used are:

XPg_1stF (5’-ACCGAACGCAACGCACGCA-3’)

XPg_1stR (5’-ACGACCACGACCGACCCGAACCACGA-3’).

A second PCR was performed on the PCR product using the following protocol: 3 minutes at 94°C followed by 12 cycles of 30 seconds 94°C, 10 seconds 66°C, and 10 seconds 72°C, followed by final extension at 72°C for 2 minutes then hold at 4°C. The sequences of the primers used are: P7_XPg_2ndF (5’-AGCAGAAGACGGCATACGAGATAGCACCGAACGCAACGCACGCA-3’) unique index reverse primer P5_XPg_2ndR (5’-AGATACGGCGACCACCGAGATCTACACGCACGACGAGACGCAGACGAAXXXXXXACGACGACCGACCCGAACCACGA-3’, X bases refers to index barcode sequence).

Second PCR products were then identified through agarose gel electrophoresis (2.5%, Lonza) at expected size of 227 bp and extracted using PureLink Quick Gel Extraction Kit (Thermo Fisher Scientific). The purified PCR product was quantified using High Sensitivity D1000 ScreenTape and Agilent 4200 TapeStation System (Agilent Technologies). Barcode representation was measured by NGS using Illumina HiSeq2000 with Seq_XPg_BC30 (5’-AGACGACCTGCTCCAGCTGCACCA-3’) as read 1 sequencing primer and RSeq-IND-XP NGS (5’-ACACGCACGACGAGACGCAGACGAA-3’) as i5 index primer.

#### Quantitative Scale

A known quantity of barcoded cells was used as normalization scale as previously described (67). In brief, the barcoded cells are infected with the same lentivirus that carries a unique barcode that are not shared in the lineage tracing library with a low Multiplicity of Infection (MOI < 0.1). Resulting infected cells are then selected by puromycin and expanded for preparing the spike-in mixture with following cells:

50 cells carrying the bc14-001.bc30-100002 barcode (CCAAAGATAGATCATGGTACACCATGACACGTCACATGTGTGCAGTCA);

500 cells carrying the bc14-001.bc30-100003 barcode (CCAAAGATAGATCATGGTACTGACACTGCAGTACCACATGCACACAAC);

5,000 cells carrying the bc14-001.bc30-100001 barcode (CCAAAGATAGATCATGGTCATGGTACACACGTGTCAACTGACCACAAC);

50,000 cells carrying the bc14-001.bc30-100004 barcode (CCAAAGATAGATCATGGTTGACACACACTGACTGACTGACTGACACCA);

500,000 cells carrying the bc14-001.bc30-100005 barcode (CCAAAGATAGATCATGGTGTACACGTACGTACTGACCAACTGACGTCA).

During the tissue and cell dissociation step of barcode library preparation, the cell mixtures were added and served as a reference for spike-in normalization of the sequencing reads. After normalizing read counts of each library by the size factor (DNA content) and library size, a high correlation coefficient (r=0.96) was observed between normalized read counts and cell number in log scale. This calibration curve was then used to predict cell count of each barcode in tumors.

### *In vivo* Experiments

#### Mouse strains

Conditional *Ptf1a*Cre;*LSL-Trp53^fl/+^*and *LSL-Kras^G12D/+^* strains were interbred to obtain *Ptf1a*Cre;*LSL-Kras^G12D/+^;LSL-Trp53^fl/fl^;*double mutant animals (hereafter KP^-/-^C mouse model) and kept in a C57BL/6 background (47). Conditional *Ptf1a*Cre; *LSL-Kras^G12D/+^* (obtained from Dr. Haoquang Ying, UTMDACC) and (46) *LSL-Trp53^R172H/+^* (obtained by Dr. Guillermina Lozano, UTMDACC) strains were interbred to obtain *Ptf1a*Cre;*LSL-Kras^G12D/+^;LSL-Trp53^R172H/+^;*double mutant animals (hereafter KPC mouse model) and kept in pure C57BL/6 background. Nude, NSG and C57BL/6 female mice (6 weeks old) were purchased from Experimental Radiation Oncology at MD Anderson Cancer Center. For treatment with anti-CD8 and anti-PD1 monoclonal antibodies, C57BL/6J female mice (6 weeks old) were purchased from Jackson Laboratory. All animal studies and procedures were approved by the UTMDACC Institutional Animal Care and Use Committee. All experiments conformed to the relevant regulatory standards and were overseen by the institutional review board. No sex bias was introduced during the generation of experimental cohorts. Littermates of the same sex were assigned randomly to experimental arms.

#### Pancreas orthotopic injections and surgical procedures

C57BL/6 female mice (8 weeks old) were shaved and anesthetized using isoflurane (Henry Schein Animal Health). Analgesia was achieved with buprenorphine SR (0.1 mg/Kg BID) (Par Parmaceutical) *via* subcutaneous injection, and shaved skin was disinfected with Chlorhexidine and 70% ethanol. A 0.5-cm incision was performed on the left flank through the skin/subcutaneous and muscular/peritoneal layers. Pancreas was exposed, and 20 μL of cell resuspension (total 1×10^5^ KPC or KP^-/-^C cells per mouse) were injected using a 27 Gauge Hamilton syringe. Hemostasis was controlled with a bipolar cautery (Bioseb) if needed. The pancreas was carefully repositioned into the abdominal cavity, and muscular/peritoneal planes were sutured individually by absorbable sutures. The skin/subcutaneous planes were closed using metal clips. Mice were monitored daily for the first three days, and twice/week thereafter for signs of tumor growth by manual palpation or MRI when appropriate.

#### Subcutaneous transplantation, tumor volume measurement and survival study

Tumor single cell suspension was resuspended in DMEM without phenol Red (Sigma) and Matrigel (BD Biosciences, 356231) (1:1 dilution) at a density of 1×10^5^ cells/100μL for KPC and KP-/-C cells and 1×10^6^ cells/100μL for PATC53 and PATC66 cells. Cell suspensions were injected subcutaneously into the flank of 8-week-old NSG, nude or C57BL/6 female mice as indicated in figures. Mice were monitored every 5 days, and tumor measurements was calculated according to the formula: Tumor volume (mm^3^) = (W^2^xL)/2, where W is Width and L is length. Mice that underwent orthotopic cell transplantation were monitored over time for tumor development by magnetic resonance imaging and survival annotated.

#### Treatments

The following antibodies were purchased from Bio X cell: InVivoMab anti-mouse PD-1 (CD279) (clone 29F.1A12, # BE0273), InVivoMab anti-mouse CD8 (clone YTS 169.2, # BP0117) and InVivoMab rat IgG2a isotype control (clone 2A3, # BE0089). Treatments with monoclonal antibody against PD-1 or CD8 or control were started 7 days after KPC cell subcutaneous injection in C57BL/6 mice. Monoclonal antibodies were diluted in PBS to a final concentration of 200μg/100μL and 50μg/100μL respectively for anti-PD-1 and anti-CD8. The treatments were performed by intraperitoneal injection of 100μL of antibody per 25g mouse, three times a week for two consecutive weeks.

#### Euthanasia, necropsy, and tissues collection

Mice were euthanized by exposure to CO_2_ followed by cervical dislocation. A necropsy form was filled in with mouse information, tumor size and weight, infiltrated organ annotations, and metastasis number and location.

#### Non-invasive MRI imaging

A 7T Bruker Biospec (BrukerBioSpin), equipped with 35mm inner diameter volume coil and 12cm inner-diameter gradients, was used for MRI imaging. A fast acquisition with relaxation enhancement sequence with 2,000/39 ms TR/TE, 256×192 matrix size, r156uM resolution, 0.75mm slice thickness, 0.25mm slice gap, 406×30cm FOV, 101kHz bandwidth, and 4 NEX was used for acquired in coronal and axial geometries a multi-slice T2-weighted images. To reduce respiratory motion, the axial scan sequences were respiratory gated. All animal imaging, preparation, and maintenance was carried out in accordance with MD Anderson’s Institutional Animal Care and Use Committee policies and procedures.

#### Cell Isolation from tumors

Tumors from mouse xenografts were harvested in HBSS (Gibco) and kept on ice. Isolation of pancreatic tumor cells was performed by mechanical dissociation (mincing the tumor tissue with sterile scissors) and enzymatic digestion 1 hr at 37°C in in gentle rotation (Dispase II 1 mg/ml, Collagenase IV 2 mg/ml in DMEM high glucose supplemented with 1% FBS). The single-cell populations were seeded at in DMEM high glucose (Gibco) supplemented with 10% FBS (Gibco), 100 IU/ml penicillin (Gibco), and 100 μg/ml streptomycin (Gibco). In order to get rid of the murine fibroblasts in the culture, we performed brief trypsinization cycle (0.25% Trypsin-EDTA, Gibco). For single-cell RNA and flow-cytometry analyses, tumors from KPC cells were minced into small fragments and digested with 1.5 mg/mL collagenase IV and 50 U/mL DNase I for 30min at 37°C under agitation. For flow cytometry analysis, the lymph nodes were collected and passed through 40 μm cell strainers and used for compensation.

### In Silico Analyses

#### Analysis of the TCGA dataset

For TCGA GSEA analysis, the TCGA pancreatic adenocarcinoma mRNA dataset is downloaded from Broad GDAC website (http://firebrowse.org/?cohort=PAAD&download_dialog=true). The mRNA data from the Illumina Genome Analyzer platform in the website was used here (PAAD.rnaseqv2 illuminahiseq_rnaseqv2 unc_edu Level_3 RSEM_genes_normalized data.data.txt). There was a total of 179 tumor samples and 4 normal adjacent tissues with their mRNA available. The clinical survival profile of 179 samples was downloaded from cbio-portal website (http://www.cbioportal.org). The tumor purity of 179 TCGA RNA samples was computed by the ABSOLUTE developed by Broad Institute. Only those sample with purity above 50 percent were selected. These 70 high-purity tumor samples were classified into high and low abundant groups for survival analysis through applying median value as the cut off. Survival was analyzed according to the Kaplan-Meier method, and differences in their distribution were evaluated by means of the log-rank test.

#### Analysis of the PRINCE dataset (70)

Median DPY30 mRNA expression from metastatic sites was used to segregate patient into high- or low-DPY30 expression and generate progression-free and overall survival Kaplan-Meier plots.

#### Automatic image segmentation and quantification

Fluorescence quantification was done using MATLAB (The MathWorks, Inc.). Specifically, Otsu’s thresholding method and marker-controlled watershed algorithm was used to segment and specify nuclear regions of individual cells. Expression of each marker in the individual segmented area was then quantified by pixel intensity.

#### ChIP-Seq data analysis

Samples were sequenced using NextSeq500 at UTMDACC Advanced Technology Genomics Core. Raw reads for each ChIP-seq were processed using Trimmomatic (v.0.39) (90) to trim off adaptor sequences incorporated in the read and to remove low quality bases, using the following parameters ‘TRAILING:3 LEADING:3 ILLUMINACLIP:adapters.fasta:2:30:10; SLIDINGWINDOW:4:15 MINLEN:50’. Then, the resulting reads were mapped to the to human genome build GRCh38/hg38 using Bowtie2 v.2 (91) with the ‘–very-sensitive’ preset of parameters. Reads that did not align to the nuclear genome or aligned to the mitochondrial genome were removed and duplicate reads were marked and removed using SAMtools (v.1.12) (92). Peak calling versus the corresponding input genomic DNA was performed using MACS2 (v.2.2.7.1) (93), for each sample separatly, with the ‘–nomodel’, ‘–extsize 200’, ‘– keep-dup all’ and ‘–qvalue 0.05’ flags and arguments. Peaks blacklisted by the ENCODE consortium analysis of artifactual signals in human cells (https://sites.google.com/site/anshulkundaje/projects/blacklists) were removed using bedtools (v.2.30) (94). To obtain a confidence peak set, only intersections between biological replicates were retained and shown as a heat map (generated using deepTools v.2.5.3) (95). Finally, differential binding analysis was performed using DiffBind (v.3.0) (96) with settings to use DESeq2 method, full library size and with subtraction of control input read counts. Binding events with an FDR cutoff of 0.05 were defined as significantly different among experimental conditions. Mean normalized reads across the three replicates obtained from DiffBind analysis were used for scatter plot representation using the ggplot2 R package (97).

#### Proteomic data analysis

Mass spectrometry data were searched using a local copy of Mascot (Matrix Science) with the following parameters: *enzyme:* Trypsin/P, *database:* SwissProt Human or Mouse (concatenated forward and reverse plus common contaminants), *fixed modifications:* Carbamidomethyl (C), *variable modifications:* Oxidation (M), Acetyl (N-term), Pyro-Glu (N-term Q), Deamidation (N,Q), *mass values:* Monoisotopic, *peptide Mass Tolerance:* 10 ppm, *fragment mass tolerance:* 0.02 Da, *Max Missed Cleavages:* 2. Mascot DAT files were parsed into Scaffold (Proteome Software) for validation, filtering and to create a non-redundant list per sample. Data were filtered using 1% protein and peptide FDR and requiring at least two unique peptides per protein.

#### ATAC-seq data analysis

Paired-end ATAC-seq fragments of human cells were aligned to mouse reference build GRCh38/hg38 using Bowtie2 v.2. (91). Reads that did not align to the nuclear genome or aligned to the mitochondrial genome were removed and duplicate reads were marked and removed using SAMtools (92). Read positions were corrected for transposon insertion offset as described (98), then only uniquely mapped paired-end reads were kept for further analysis. All ATAC-seq peaks that were no more than 3 kb away from annotated gene TSS (GENCODE v19) were selected as promoter (proximal) regions, using annotatePeaks script from HOMER package (99).

#### Whole exome sequencing data analysis

DNA sequencing of samples was performed on the Illumina HiSeq2000 sequencing platform at the UTMDACC Advanced Technology and Genomics Core. Sequencing fasta files were applied to FastQC (100) for quality control, adapters were trimmed by trimmomatic (90), and the genomic fragments were aligned to mm10 using bwa mem (101), then sorted and indexed by samtools (92). Following Base Quality Score Recalibration (BQSR) by GATK(102). Somatic mutations were called by Mutect2 and copy number segments were called by CNVkit (103).

#### RNA-Seq data analysis

RNA sequencing of samples was performed on the Illumina NovaSeq6000 sequencing platform at the UTMDACC Advanced Technology and Genomics Core. FastQC v0.11.9 (https://github.com/s-andrews/FastQC) was used to perform quality control. Raw reads were first aligned to the human genome GRCh38.p14 or mouse genome GRCm38.p6 using STAR2.7.10b v3.0.0 (104) and sorted using samtools v1.16 (92). Gene expression was quantified using Rsubread v2.14.2 (105). Differential expression analysis was performed using edgeR v3.42.4 (106) and limma v3.56.2 (107). Pathway enrichment analysis was performed using R package fgsea v1.26.0, clusterProfiler v4.8.2, msigdbr v7.5.1, ReactomePA v1.44.0 (108).

#### Single-cell RNA-Seq analysis

Tumors deriving from KPC cells were digested and single cell suspension was diluted in 2% FBS in PBS for further processing. Chromium single-cell sequencing technology from 106× Genomics was used to perform single-cell separation, complementary DNA amplification and library construction following the manufacturer’s guidelines. Briefly, the cellular suspensions were loaded on a 10x Chromium Single Cell Controller to generate single-cell gel bead-in-emulsions. The scRNA-seq libraries were constructed using the Chromium Single Cell 3ʹ Library & Gel Bead Kit v.2 (PN-120237, 10x Genomics). The HS dsDNA Qubit Kit was used to determine the concentrations of both the cDNA and the libraries. The HS DNA Bioanalyzer was used for quality-tracking purposes and size determination for cDNA and lower-concentrated libraries. Sample libraries were normalized to 7.5 nM and equal volumes were added of each library for pooling. The concentration of the library pool was determined using the Library Quantification qPCR Kit (KAPA Biosystems) before sequencing. The barcoded library at the concentration of 275 pM was sequenced on the NovaSeq6000 (Illumina) S2 flow cell (100 cycle kit) using a 26 × 91 run format with 8 bp index (read 1). To minimize batch effects, the libraries were constructed using the same versions of reagent kits and following the same protocols, and the libraries were sequenced on the same NovaSeq6000 flow cell and analyzed together.

The raw scRNA-seq data were pre-processed (demultiplex cellular barcodes, read alignment, and generation of gene count matrix) using Cell Ranger Single Cell Software Suite (v7.1.0,10x Genomics). Scrublet v0.2.3 (109), an algorithm to predict doublets in scRNA-seq data, was used to further clean doublets. Seurat (version 5.0.2) (110) was applied to the normalized gene-cell matrix to identify highly variable genes (HVGs) for unsupervised cell clustering. Principal component analysis (PCA) was performed on the top 2000 HVGs. The FindNeighbors function of Seurat was used to construct the Shared Nearest Neighbor (SNN) Graph, based on unsupervised clustering performed with Seurat function FindClusters and top 60 PCs. Different resolution parameters for unsupervised clustering were then examined, and cluster marker genes were checked to determine the optimal number of clusters with distinct transcriptional profiles. For visualization, the dimensionality was further reduced using Uniform Manifold Approximation and Projection (UMAP) method (111) with Seurat function RunUMAP. The PCs used to calculate the embedding were the same as those used for clustering. Differentially expressed genes (DEGs) were identified for each cluster using the FindMarkers function in Seurat (110). The top-ranked DEGs and the enrichment of canonical marker genes were integrated, together with global cluster distribution and prior knowledge to infer major cell types and cellular states.

#### Re-replication sequencing

Briefly, 30×10^6^ control or Dpy30-KO KPC cells were seeded in a square bioassay dish (Corning®) and were synchronized in S phase using a double thymidine block ad described above. After synchronization, cells were released in complete medium supplemented with 25 μM CldU for 45 minutes. Cells were then washed twice with PBS and a second pulse in complete medium supplemented with 250 μM IdU was added and incubated for 45 minutes. Cells were then collected, and cell pellets were lysed in 350 μL of lyses buffer (12mM Trish-HCl, 0.1x PBS, 6 mM EDTA, 0.5% SDS) supplemented with 40 μL Proteinase k (20mg/ml) and 10 μL RNAse A (10mg/ml) for 10 minutes at RT and then 15 minutes at 56°C. Samples were sonicated to obtain chromatin fragments of ∼200 - 600 base pairs (bp). After collecting the 10% of the sample as input control, samples were diluted using 1.6 mL of ChIP dilution buffer (10 mM Tris-HCl, 140 mM NaCl, 0.1% DOC, 1% Triton-X, 1 mM EDTA). Diluted DNA samples were then denatured for 10 min at 95°C and cool down at 4°C to preserve ssDNA. Next, ssDNA samples were immunoprecipitated as described above using Dynabeads Protein G (Invitrogen) pre-conjugated with antibodies against CldU (rat monoclonal anti-BrdU clone BU1/75) and IdU (mouse monoclonal anti-BrdU clone B44). Eluted ssDNA samples and inputs were purified with PCR Clean UP kit and eluted in water. ChIP DNA was prepared with xGen ssDNA & low input DNA library preparation kit and sequenced using NovaSeq X Plus 10 Billion 26×150 PE (142M PE reads per sample).

The raw fastq files were processed using a pipeline of Nextflow. In short, the raw reads were trimmed to remove adapter sequences, aligned to mouse mm10 genome, and filtered as described (54). The filtered alignments were then merged, and the CldU/IdU ratios were calculated with deepTools and normalized with the default window size of 5000 and loess span of 300000. The coverage tracts of individual and merged bam files were generated with deepTools (v3.5.4).

## Statistical Analysis

*In vitro* and *in vivo* data are presented as the median values or mean ± S.D. (standard deviation). Statistical analyses were performed using Student’s t-test, one-way or two-way ANOVA test as specified in figure legends. Results from survival experiments were analyzed with a Log-rank test and expressed as Kaplan–Meier survival curves.

## Acknowledgments

We thank Michael Kim for PDX-derived PDAC cell lines; all Draetta lab members and Guoxin Feng for scientific discussion; R. Nguyen and J. Martinez in the Department of Genomic Medicine at MD Anderson for lab management; Michael Peoples and Vandhana Ramamoorthy with the MD Anderson TRACTION platform for valuable support with cloning; Alejandro Hernandez Martinez, Edward Q. Chang and Ningping Feng with the MD Anderson TRACTION platform for support with tissue imaging acquisition and drug formulation; and Vivien Van, Charles Kingsley and Jorge Delacerda at the MD Anderson Small Animals Imaging Facility; The Animal Support Facility (RASF), which was partially funded by the MD Anderson Support Grant CA016672; the MD Anderson Department of Veterinary Medicine; the MD Anderson Advanced Microscopy Core, which was funded by NIH S10 RR029552; Asha Multani and the MD Anderson Cytogenetics and Cell Authentication Core; Erika Thompson, Hongli Tang, David Pollock and the MD Anderson Advanced Technology Genomics Core (ATGC), which was funded by the CA016672 Core Grant; Karen Ramirez, David Dwyer, Veena Papanna, and the MD Anderson Advanced Cytometry and Sorting Facility, which was funded by the NCI Cancer Center Support Grant P30CA16672; and Genscript for support and service. The mEmerald-MCM vectors were a gift from Michael Davidson, the lentiCRISPR v2 vector was a gift from Feng Zhang. The pcDNA3.1 Flag-hDPY30 vector was a gift from Kai Ge. The psPAX2 and pMD2G vectors were a gift from Didier Trono. Schematics were created using BioRender.com. This research was supported by the 2022 AACR−AstraZeneca START Grant, Grant Number 22-40-12-CITR to F.C.; AIRC and the European Union’s Horizon 2020 research and innovation program under the Marie Skłodowska-Curie grant agreement no. 800924 to F.C.; CPRIT Training Award (RP210028) to E.Y.; N.P. is a TRIUMPH Fellow in the CPRIT Training Program (RP210028); Cancer Research and Prevention Institute of Texas (CPRIT) grant RP190599, NIH/NCI R01 CA258917, the Andrew Sabin Family Foundation grant, and the V Foundation grant (V2020-018) to A.V.; NIH/NCI SPORE in gastrointestinal cancer grant P50CA221707 to G.F.D and A.V.; UT MD Anderson Cancer Center Start-Up Funds to A.V.; UT MD Anderson Cancer Center Pancreatic Cancer Moon Shot and Pancreatic Cancer Action Network (PanCAN) to G.F.D.; NIH/NCI P01 CA117969-12 to A.V. and G.F.D. G.F.D. was also supported by the Sewell Family Distinguished University Chair in Genomic Medicine.

## Author contributions

Conceptualization: F.C., A.V., G.F.D.; Methodology: F.C., L.P., I.H., C.B., E.Y., Z.L., L.Z., L.C.M, P.J., S.A., M.D.P., W.Y., A.Z., C.J.; Software: Y.C., C.B., L.Z., Z.L., E.Y., S.S., L.W., G.T., C.Bo.; Validation: F.C., L.P, I.H, E.Y., L.C, H.K., R.S., Z.C., I.S., N.P., S.A., K.C.C, C.A.D., C.L., M.M., K.R., C.J.; Formal Analysis: F.C., L.P., Z.L., C.B., L.Z., E.Y., S.S., M.D.P.; Investigation: F.C., L.P, I.H, E.Y., K.C.C., L.C.M., M.D.P.; Resources: F.C., L.P., I.H., C.B., Z.L., L.Z., E.Y., R.S., L.C.M., S.A., S.J., B.G.H., B.V., C.L., M.M., K.R., H.W., W.Y., V.G., T.H., G.T., L.W., M.D.P.; Data curation: F.C., L.P., I.H., Z.L., C.B., L.Z., E.Y.; Writing – Original draft: F.C., A.V., G.F.D.; Writing – Review and Editing: F.C., A.V., G.F.D., S.G.; Visualization: F.C., C.B., Z.L., E.Y.; Supervision: F.C., A.V., G.F.D; Project Administration: F.C., A.V., G.F.D; Funding Acquisition: F.C., A.V., G.F.D;

---

**Extended Data Figure 1.**
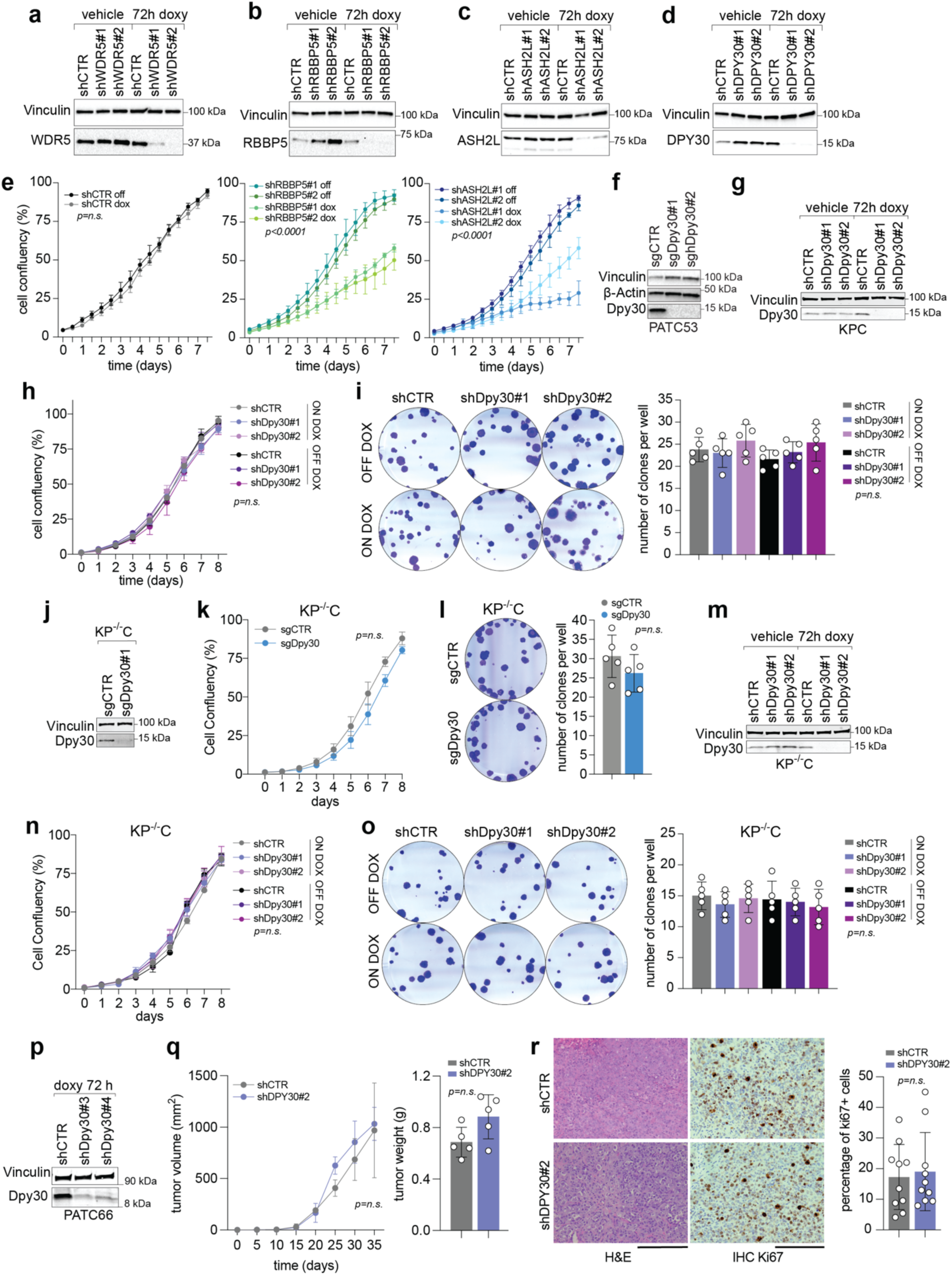
DPY30 is dispensable for cell proliferation. **a-d.** Western Blot analysis of PATC53 cells harboring shRNAs against WDR5, RBBP5, ASH2L, DPY30 or control (respectively shWDR5, shRBBP5, shASH2L, shDPY30 or shCTR). **e.** Growth curve analysis of PATC53 cells described in a-d. **f.** Western Blot analysis of control (sgCTR) or DPY30-KO PATC53 cells. **g.** Western Blot analysis of KPC cells silenced for Dpy30 (shDPY30) or control (shCTR). **h.** Growth curve of KPC cells described in g. **i.** Representation (left) and quantification of a clonogenic assay of KPC cells, as described in g. **j.** Western Blot analysis of sgCTR or sgDpy30 KP^-/-^C cells. **k.** Growth curve of KPC cells described in j. **l.** Representation (left) and quantification of a clonogenic assay of KP^-/-^C cells, as described in j. **m.** Western Blot analysis of shCTR or shDpy30 KP^-/-^C cells. **n.** Growth curve of KP^-/-^C cells described in m. **o.** Representation (left) and quantification (right) of a clonogenic assay of KP^-/-^C cells, as described in m. **p.** Western Blot analysis of shCTR or shDPY30 PATC66 cells. **q.** Volume (left) and weight (right) of tumors from shCTR or shDPY30 PATC66 cells injected subcutaneously into the flank of NSG mice. **r.** Histopathological analysis of tumors described in (q). Hematoxylin and eosin (H&E) staining (left), Ki67 IHC staining (middle) and Ki67 quantification (right) are shown. In (a-e, g-I, m-r) cells were treated or not with doxycycline to induce shRNA expression. In western blot analyses vinculin or actin were used as loading controls. In e, h,i, k, l, n, o, q, r data are expressed as mean ±SD. Each dot represents a biological replicate. Statistical significance was calculated using one-way ANOVA followed by Tukey’s multiple comparison tests (e, h, i, n, o) or using Student’s t test (k, l, q, r).

**Extended Data Figure 2.**
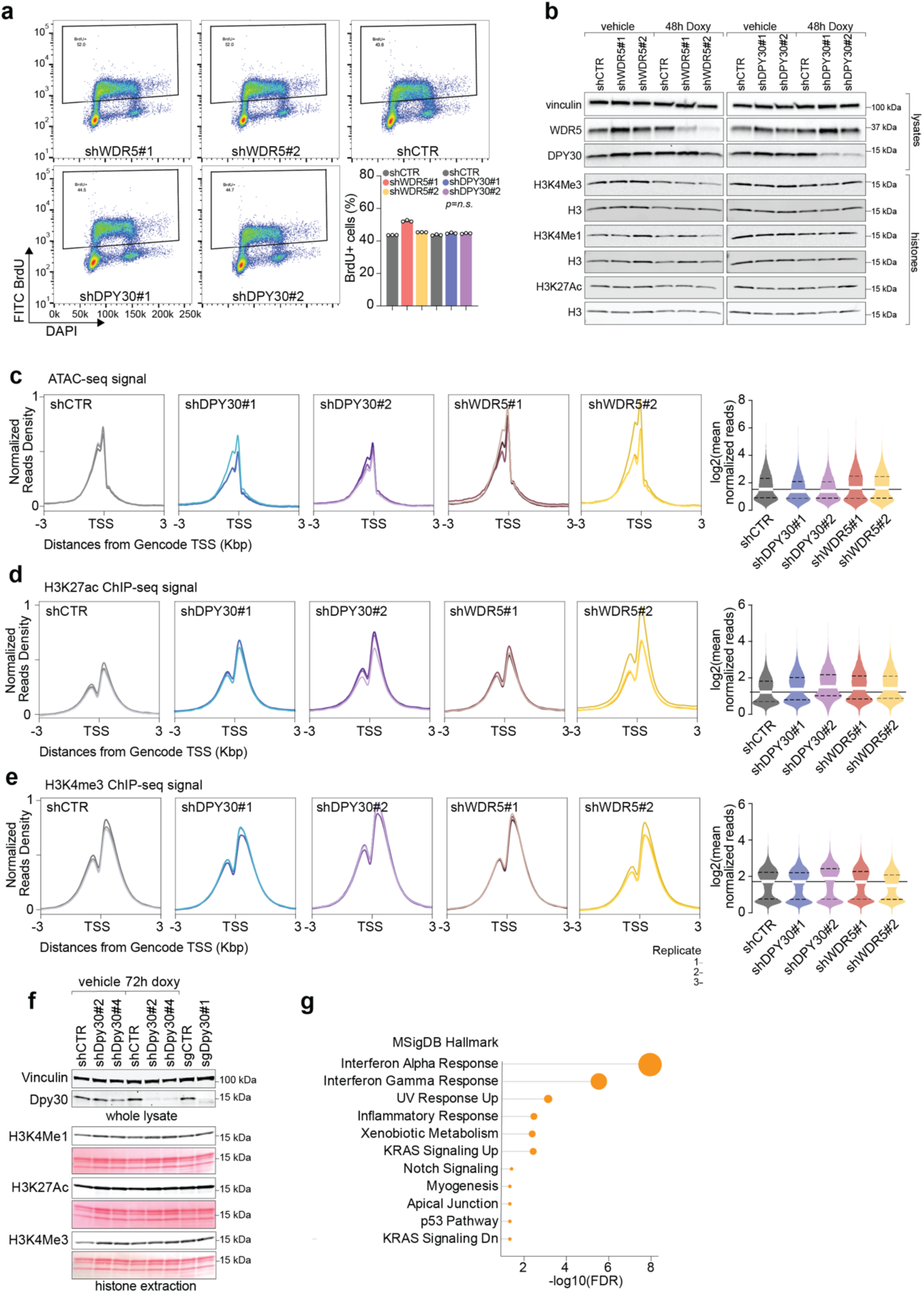
WDR5 loss, but not DPY30, broadly affect H3K4 methylation patterns. **a.** Dot plots of BrdU incorporation assay of PATC53 cells silenced for WDR5 or DPY30 (respectively shWDR5 and shDPY30) or control (shCTR). Cells were treated with doxycycline for 48 hrs to induce shRNA expression. Data are expressed as mean ±SD, each dot represents a biological replicate. Statistical significance was calculated using one-way ANOVA. **b.** Western Blot analysis of whole lysate (upper panel) or histone lysates (lower panel) in PATC53 cells, as described in a. **c.** Plots showing normalized reads intensity (left) and the quantification (right) of ATACseq signals from three biological replicates of PATC53 cells as described in a-b. **d.** Plots showing normalized reads intensity (left) and the quantification (right) of H3K27ac signals from three biological replicates of PATC53 cells in a-b. **e.** Plots showing normalized reads intensity (left) and the quantification (right) of H3K4me3 signals from three biological replicates of PATC53 cells as described in a-b. **f.** Western Blot analysis of whole lysate (upper) or histone lysates (lower) in KPC cells silenced/knock out for Dpy30 (respectively shDpy30 and sgDpy30) or control (shCTR or sgCTR). Cells were treated with doxycycline or vehicle for 72 hrs to induce shRNA expression. **g.** Plot representing the top regulated pathways resulting from genes having H3K4me3 peak loss in shWDR5 PATC53 cells compared to controls (shCTR). In c-d, data in the violin plots are presented as median value.

**Extended Data Figure 3.**
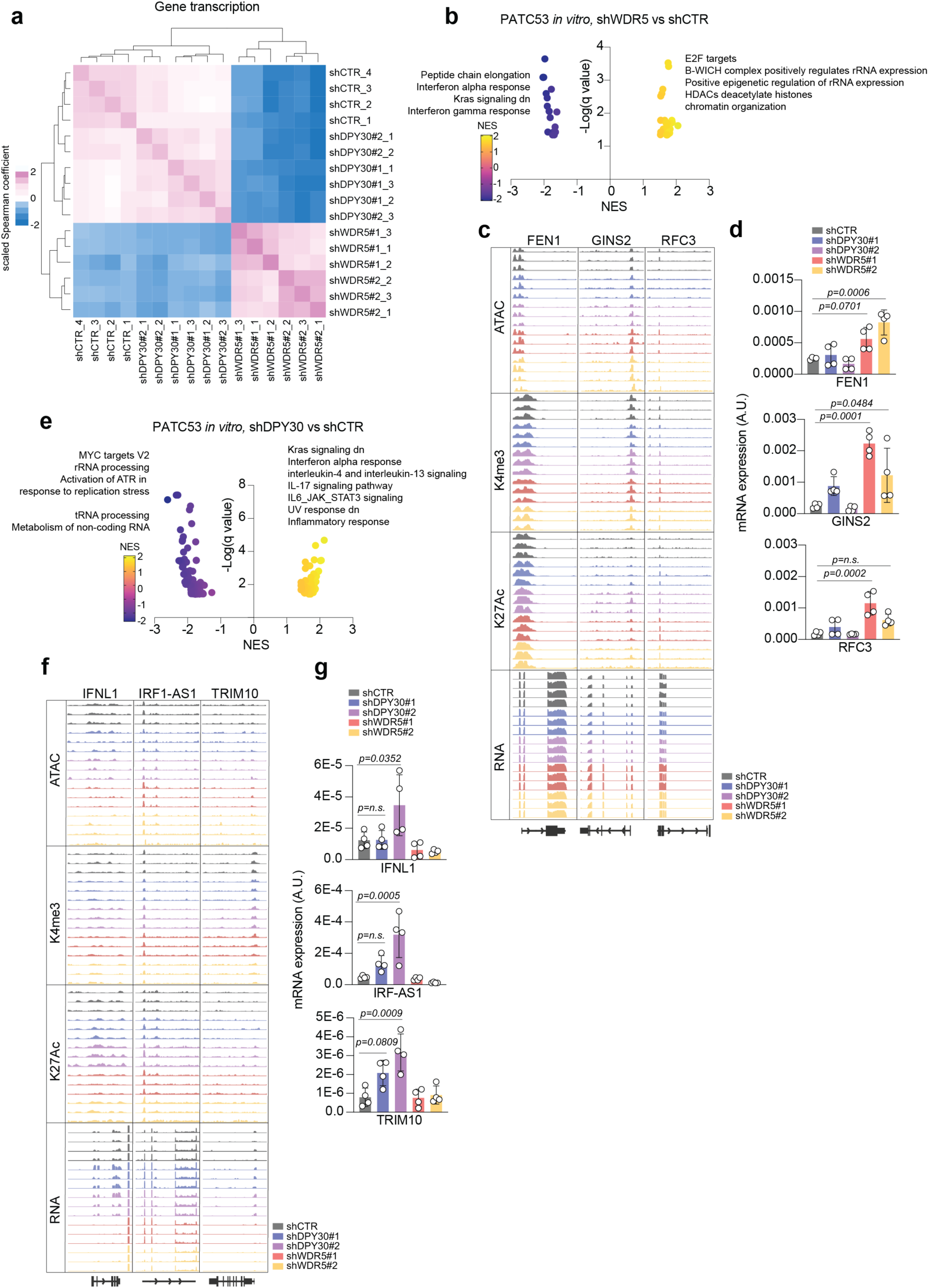
DPY30 loss moderately upregulates inflammatory pathways. **a.** Heatmap showing Spearman correlation between samples and biological replicates for bulk RNA sequencing (RNAseq) of PATC53 cells silenced for WDR5 (shWDR5) or DPY30 (shDPY30) or control (shCTR). **b.** Volcano plot representing the GSEA results from bulk RNAseq data in PATC53 cells shWDR5 versus shCTR. **c.** Impact of shCTR, shDPY30 or shWDR5 in PATC53 cells on *FEN1*, *GINS* and *RFC3* analyzed by chromatin accessibility (ATAC), histone H3 methylation (H3K4me3), histone H3 acetylation (H3K27ac) and mRNA expression (RNA). **d.** Graphs reporting the mRNA expression of *FEN1*, *GINS* and *RFC3* analyzed by quantitative PCR in shCTR, shDPY30 and shWDR5 PATC53 cells. **e.** Volcano plot representing GSEA from bulk RNAseq data in PATC53 cells shDPY30 versus shCTR. **f.** Impact of shCTR, shDPY30 or shWDR5 in PATC53 cells on *IFNL1*, *IRF-AS1* and *TRIM10* analyzed by ATAC, H3K4me3, H3K27ac and RNA expression. **g.** Graphs reporting the mRNA expression of *IFNL1*, *IRF-AS1* and *TRIM10* analyzed as described in d. In b-c, top upregulated pathways are indicated. In d and g, data are expressed as mean ±SD, each dot represents a biological replicate, and statistical significance was calculated using one-way ANOVA.

**Extended Data Figure 4.**
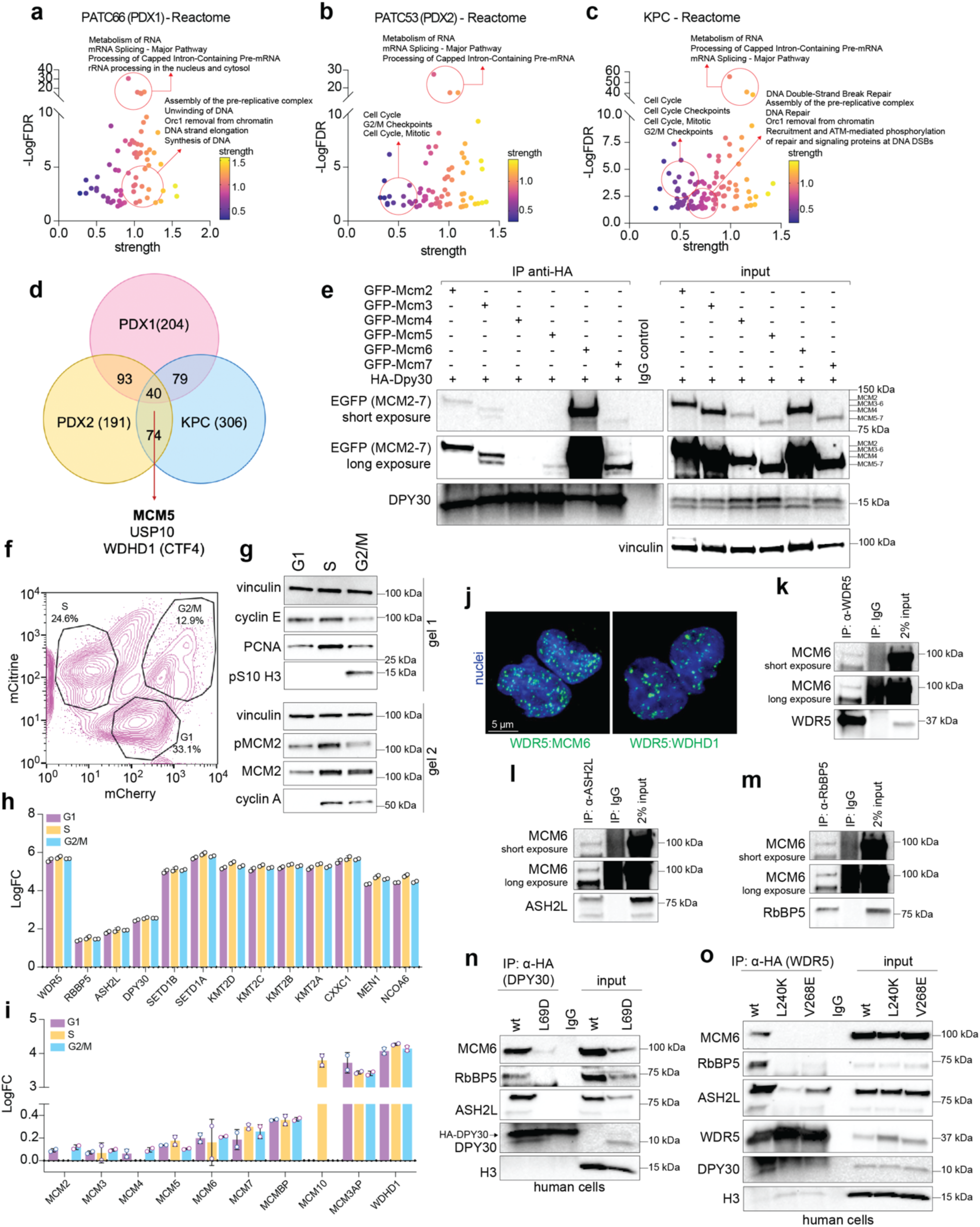
The WRAD core interacts with the replisome machinery in a non-canonical manner. **a-c.** Dot plots reporting pathway enrichment analyses of DPY30 interactors in human PATC66 (a), PATC53 (b), or mouse KPC (c) cells. Pathway enrichment is calculated using STRING online tool and Reactome module. **d.** Venn diagram showing the DPY30 common interactors among PATC53 (PDX2), PATC66 (PDX1), and KPC cells. **e.** Western Blot analysis of 293T cells co-overexpressing HA-tagged mouse DPY30 and EGFP-tagged mouse MCM members (MCM2-7). Control IgG and input were included. **f.** Contour plot showing the gating strategy for sorting PATC53 cells expressing FUCCI according to cell cycle phases. **g.** Western blot analysis of PATC53 cells sorted as in f. **h-i.** Bar plots showing the fold change enrichment of the epigenetic interactors (h) or replisome interactors (i) in DPY30 immunoprecipitates analyzed by mass spectrometry. Each dot represents a biological replicates and data are expressed as mean ± SD. **j.** Representative PLA staining detecting WDR5 interaction with WDHD1 and MCM6 in PATC66 cells. **k-m.** Western Blot analysis of the immunoprecipitation assay for endogenous WDR5 (k), Ash2L (l) and RbBP5 (m) in PATC53 cells. Control IgG and input were included. **n.** Western blot analysis of PATC53 cells expressing DPY30 wild type (WT) or mutant (L69D) isoforms. **o.** Western blot analysis of PATC53 cells expressing WDR5 wild type (WT) or mutant (L240K, V268E) isoforms. In (n, o), control IgG and input are shown, and H3 was used as a loading control.

**Extended Data Figure 5.**
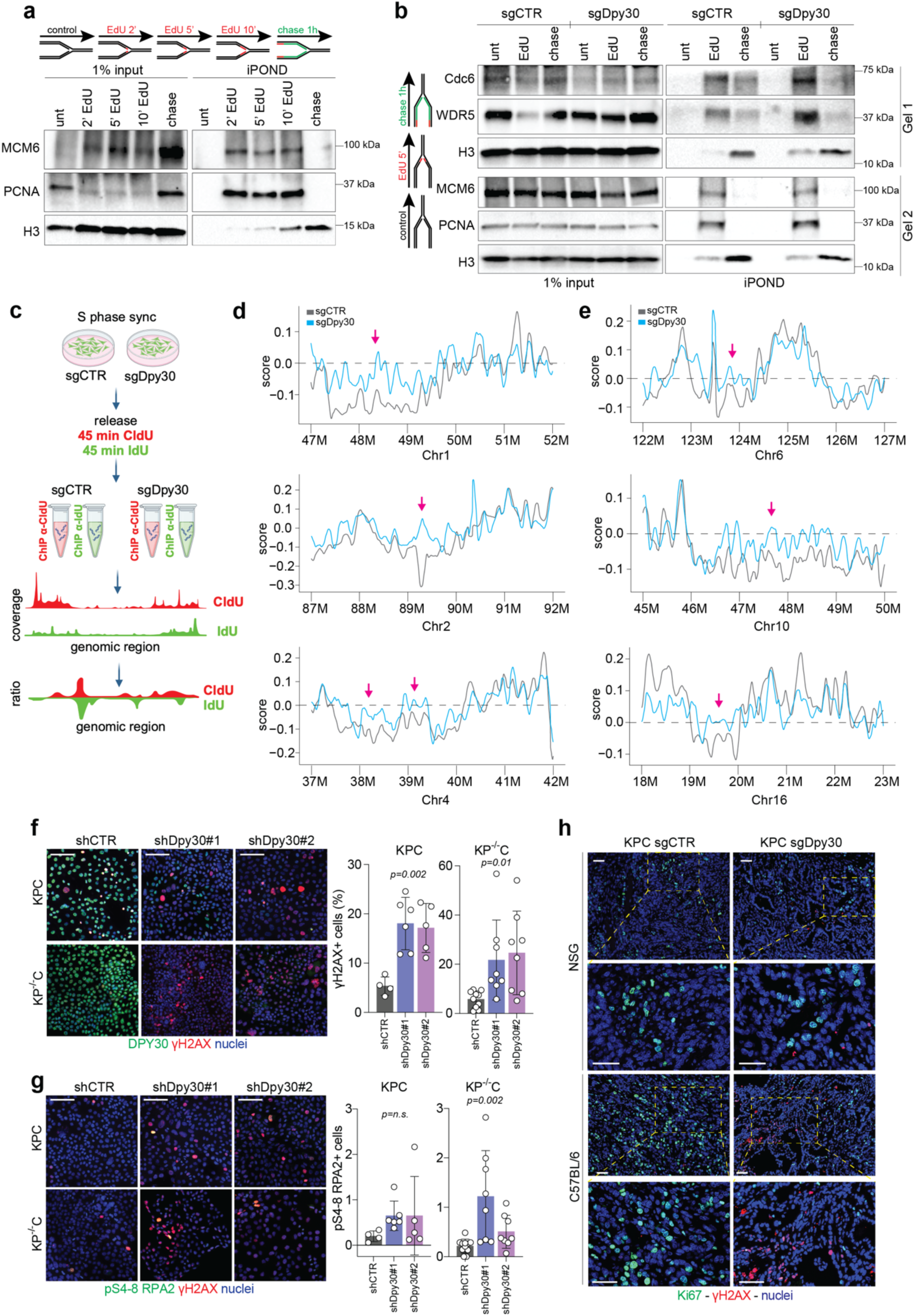
DPY30 loss leads to DNA replication stress and DNA damage. **a.** Western blot analysis of proteins bound to nascent DNA retrieved using iPOND or input control in KPC cells. Cells were treated as follows: control (negative control, no biotin in the click reaction), EdU pulse (EdU treatment for 2, 5 or 10min), chase (EdU treatment for 10 min, followed by a 60 min thymidine chase). Histone H3 was used as a loading control. **b.** Western blot analysis of iPOND or input control in KPC cells, control (sgCTR) or *Dpy30*-KO (sgDpy30). Cells were treated as described in a. Histone H3 was used as a loading control. **c.** Schematic representation of the re-replication sequencing (re-repli-seq) in KPC sg/ctr or sgDpy30 cells. **d-e.** Plots showing the genomic regions and the ratio (score) of incorporated CldU and IdU in KPC sgCTR and sgDpy30 cells. **f.** Left, panels show immunofluorescence images of KPC and KP^-/-^C cells silenced for Dpy30 (shDpy30) or control (shCTR), treated with doxycycline for 96 hrs to induce shRNA expression. In green DPY30, in red pS139 H2AX (γH2AX), in blue nuclei (DAPI). Right, graphs report the percentage of γH2AX+ in KPC cells. **g.** Left, panels show immunofluorescence staining of KPC and KP^-/-^C cells, as described in f, stained for pS4-8 RPA (green), γH2AX (red) and nuclei (blue). Right, graphs report the percentage of pS4-8 RPA+ in each cell group. **h.** Immunofluorescence staining of sgDpy30 or sgCTR KPC tumors grown in NSG (upper) or C57BL/6 (lower) mice and then stained for Ki67 (green), γH2AX (red), and nuclei (blue). In (f-g), data are expressed as mean ± SD and each dot represents a different analyzed field. Statistical significance was calculated using one-way ANOVA (f, g).

**Extended Data Figure 6.**
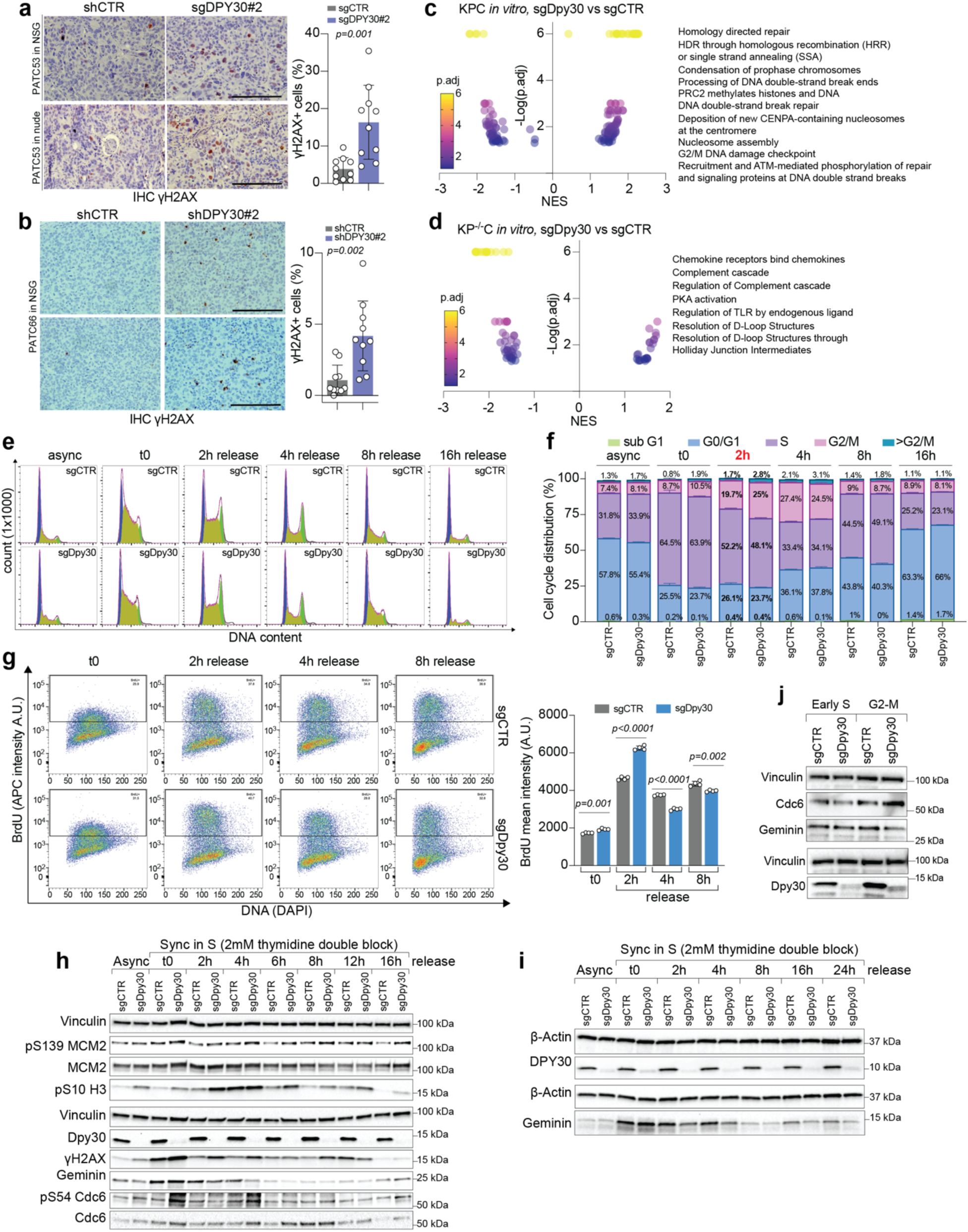
DPY30 loss leads to DNA damage and accelerated S-to-G2/M transition. **a.** Immunohistochemical staining of γH2AX (left) and quantification (right) in control (sgCTR) or DPY30-KO (sgDPY30) PATC53 tumors grown in NSG (upper) or nude (lower) mice. **b.** Immunohistochemical staining of γH2AX (left) and quantification (right) in control (shCTR) or DPY30-silenced (shDPY30) PATC66 tumors grown in NSG mice. **c-d.** Volcano plots representing the GSEA results from bulk RNAseq data in KPC (c) or KP^-/-^ C (d) cells sgDpy30 versus sgCTR. **e.** Histograms showing DNA content in sgCTR or sgDpy30 KPC cells. Cells were collected in exponential growth (async) or synchronized (t0) in S phase using a double thymidine block and then released in complete medium for the indicated time. **f.** Bar graph showing cell cycle distribution in KPC cells as described in e. **g.** Dot plots (left) and quantification (right) of the BrdU incorporation assay in KPC cells described as in e. BrdU (APC) mean intensity is shown (A.U. = arbitrary unit). **h-i.** Western Blot analysis of sgCTR or sgDpy30 KPC (h) or KP^-/-^C (i) cells, as described in e. Vinculin was used as loading control. **j.** Western Blot analysis of sgCTR or sgDpy30 KPC cells synchronized in S or G2/M phase. Vinculin was used as loading control. In (a, b, f, g), data are expressed as mean ±SD, each dot represents a biological replicate. Statistical significance was calculated using one-way student t test (a-b, g) or two-way ANOVA test (f).

**Extended Data Figure 7.**
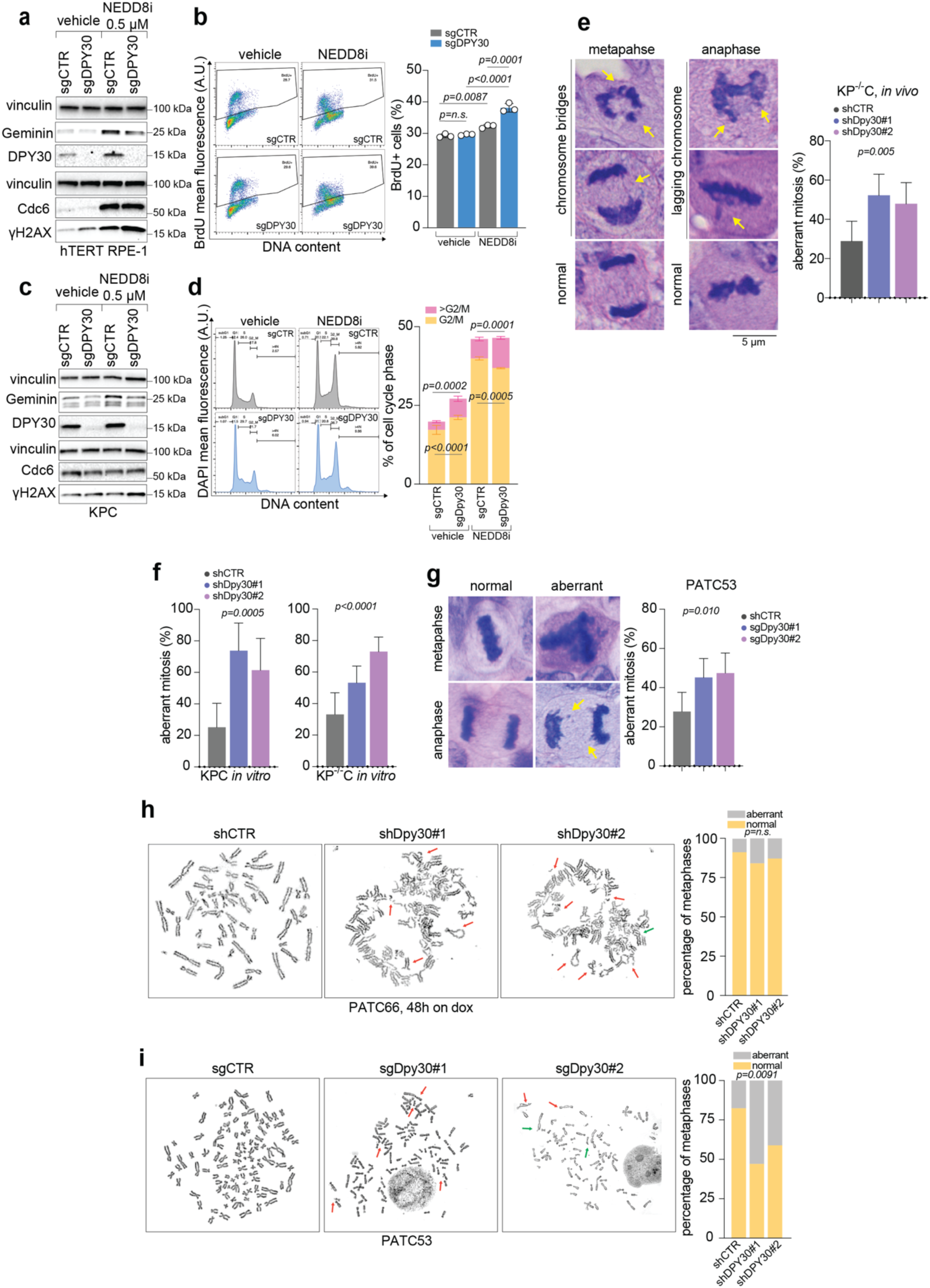
DPY30 loss increases chromosomal instability in PDAC. **a.** Western Blot analysis of control (sgCTR) or *DPY30*-KO (sgDPY30) RPE-1 cells treated with vehicle or MLN4924 for 24 hrs. **b.** Dot plots (left) and cell percentage (right) of BrdU incorporation in RPE-1 cells as described in a. **c.** Western Blot analysis of sgCTR and sgDpy30 KPC cells treated with vehicle or MLN4924 for 24 hrs. **d.** Histograms of DNA content (left) and quantification of G2/M and aneuploid (>4N) cells (right) in KPC cells, as described in c. **e. f.** Bar graphs reporting the mean percentage of aberrant mitosis in control (shCTR) or Dpy30-silenced (shDpy30) KPC and KP^-/-^C cells *in vitro*. Cells were treated with doxycycline (dox) for 96 hrs to induce shRNA expression. **g.** Left, hematoxylin and eosin (H&E) staining of mitotic sgCTR and sgDPY30 PATC53 cells grown in NSG mice. Yellow arrows indicate chromosomal laggings. Right, bar graph reporting the percentage of aberrant mitosis in PATC53 cells. **h.** Left, images of metaphasic spread in shCTR or shDPY30 PATC66 cells treated with dox for 48 hrs to induce shRNA expression. **i.** Left, images of metaphasic spread in sgCTR and sgDPY30 PATC53 cells. Right, graph reporting the percentage of normal (yellow) or aberrant (grey) mitosis. In (a, c), vinculin was used as loading control. In (h-i), red arrows indicate chromosome fusions or breaks. In (b, d-g), data are expressed as mean ±SD, dots indicate biological replicates and statistical significance was calculated using one-way ANOVA (b, e, f, g), two-way ANOVA (d) or Chi-square test (h, i).

**Extended Data Figure 8.**
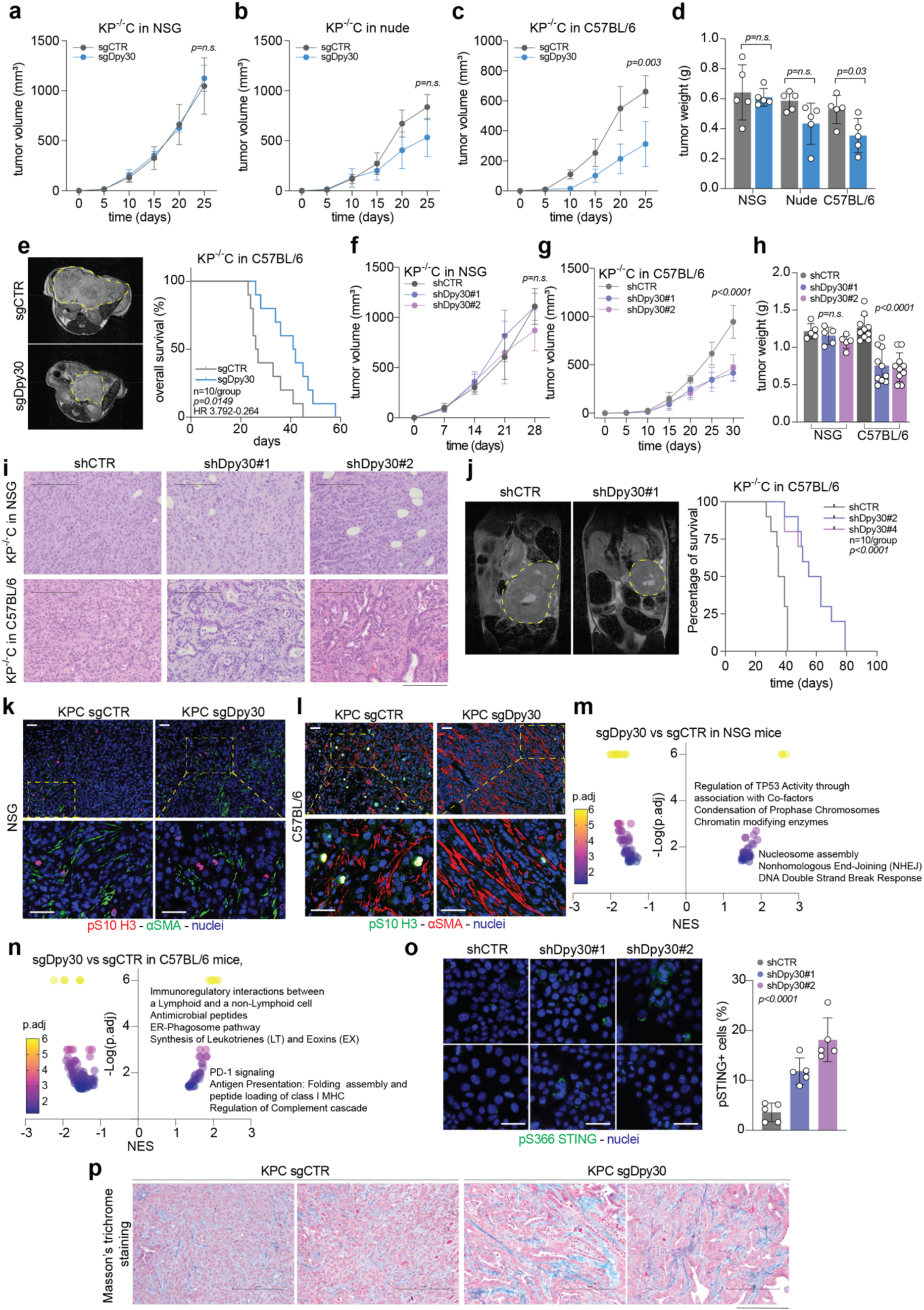
In pancreatic ductal carcinoma, DPY30 loss has a context-dependent phenotype. **a-c.** Tumor volume of control (sgCTR) and Dpy30-KO (sgDpy30) KP^-/-^C cells injected subcutaneously into the flank of NSG (a), nude (b), or C57BL/6 (c) mice. **d.** Weight (mg) of tumors described in a-c. **e.** Left, T2 Magnetic Resonance Imaging (MRI) scans of sgCTR and sgDpy30 KP^-/-^C tumors grown orthotopically in the pancreas of C57BL/6 mice. Right, Kaplan-Meier survival curve of tumor-bearing mice. **f-h.** Tumor volume (f, g) and weight (h) of control (shCTR) or Dpy30-silenced (shDpy30) KP^-/-^C cells injected subcutaneously into the flank of NSG (f) and C57BL/6 (g) mice. **i.** Hematoxylin and eosin (H&E) staining of KP^-/-^C tumors as described in f-h. **j.** Left, T2 Magnetic Resonance Imaging (MRI) scans of shCTR or shDpy30 KP^-/-^C tumors grown orthotopically in the pancreas of C57BL/6 mice. Right, Kaplan-Meier survival curve of tumor-bearing mice. **k-l.** Immunofluorescence staining of shCTR and shDpy30 KPC tumors grown in NSG (k) or C57BL/6 (l) mice and then stained for αSMA, pS10-H3, and nuclei. **m-n.** Volcano plots representing GSEA analyses from bulk RNAseq data in sgDpy30 versus sgCTR KP^-/-^C tumors grown in NSG (k) or C57BL/6 (l) mice. **o.** Immunofluorescence staining of pS366 STING (green) and nuclei (blue) in sgCTR and sgDpy30 KPC cells, in vitro. **p.** Masson’s trichrome staining of sgCTR and sgDpy30 KPC tumors grown in C57BL/6 mice. In (a-d, f-h, o), data are expressed as mean value ±SD. Dots represent biological replicates. In (e, j), statistical significance and Hazard Ratio (HR) were calculated using Log-rank test. Statistical significance was calculated using Student t test (a-d) or one-way ANOVA (f-h, o). Scale bar = 200 μm (i, p), 100 μm (k, l), 50 μm (o).

**Extended Data Figure 9.**
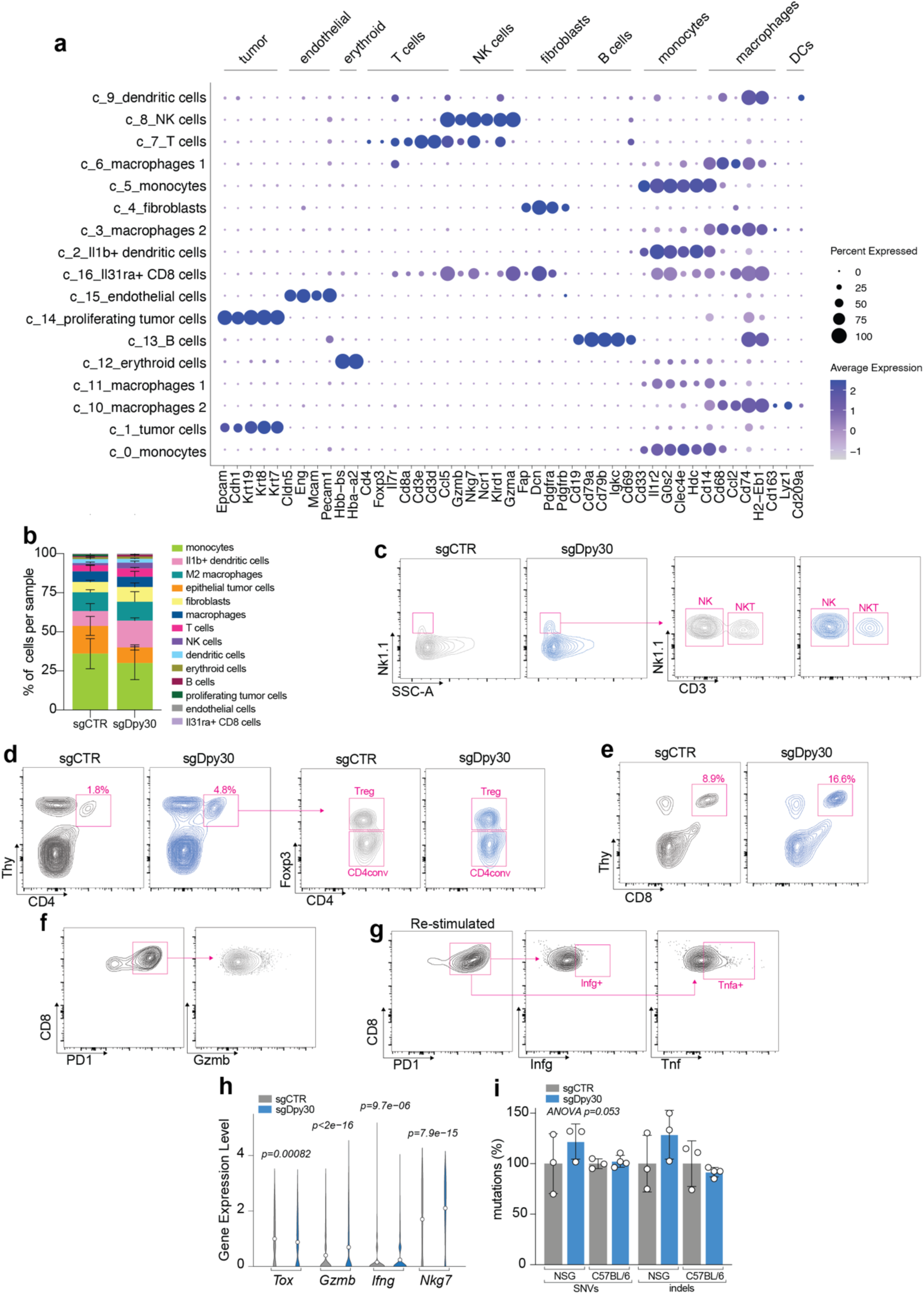
Dpy30 knockout tumors display higher T cell infiltration and activation. **a.** Bubble plot showing expression levels of genes selected for cluster annotation in single-cell RNA sequencing data from control (sgCTR) or Dpy30-KO (sgDpy30) KPC tumors grown in C57BL/6 mice. Size of dots represents the percentage of cells expressing the gene; color scale shows the average expression level. **b.** Bar plot showing the percentage of cell distribution in the different clusters (cell type) calculated using Seurat in sgCTR or sgDpy30 KPC tumors (n=3 per condition). **c-g.** Contour plots showing the flow cytometry gating strategy for detecting the NK or CD3+ NK cells (c), CD4+ conventional or Treg cells (d), CD8+ T cells (e), CD8/PD-1+ cells expressing Granzyme B (Gzmb, f), Interferon gamma (Infg, g) or Tumor Necrosis Factor (tnf, g). **h.** Violin plot showing the expression of T cell exhaustion marker (Tox) or cytotoxic markers *(Gzmb, Ifng, Nkg7)* in the T cell of sgCTR or sgDpy30 KPC tumors, calculated using Seurat. **i.** Bar plot showing the percentage of single nucleotide variants (SNVs) and indels in sgCTR or sgDpy30 KP-/-C tumors grown in NSG or C57BL/6 mice (n=3 or 4 tumors per group). Data are expressed as median (h) or mean value ±S.D. (i). Statistical significance was calculated using Student’s t test (h) or a two-way ANOVA (i).

**Extended Data Figure 10.**
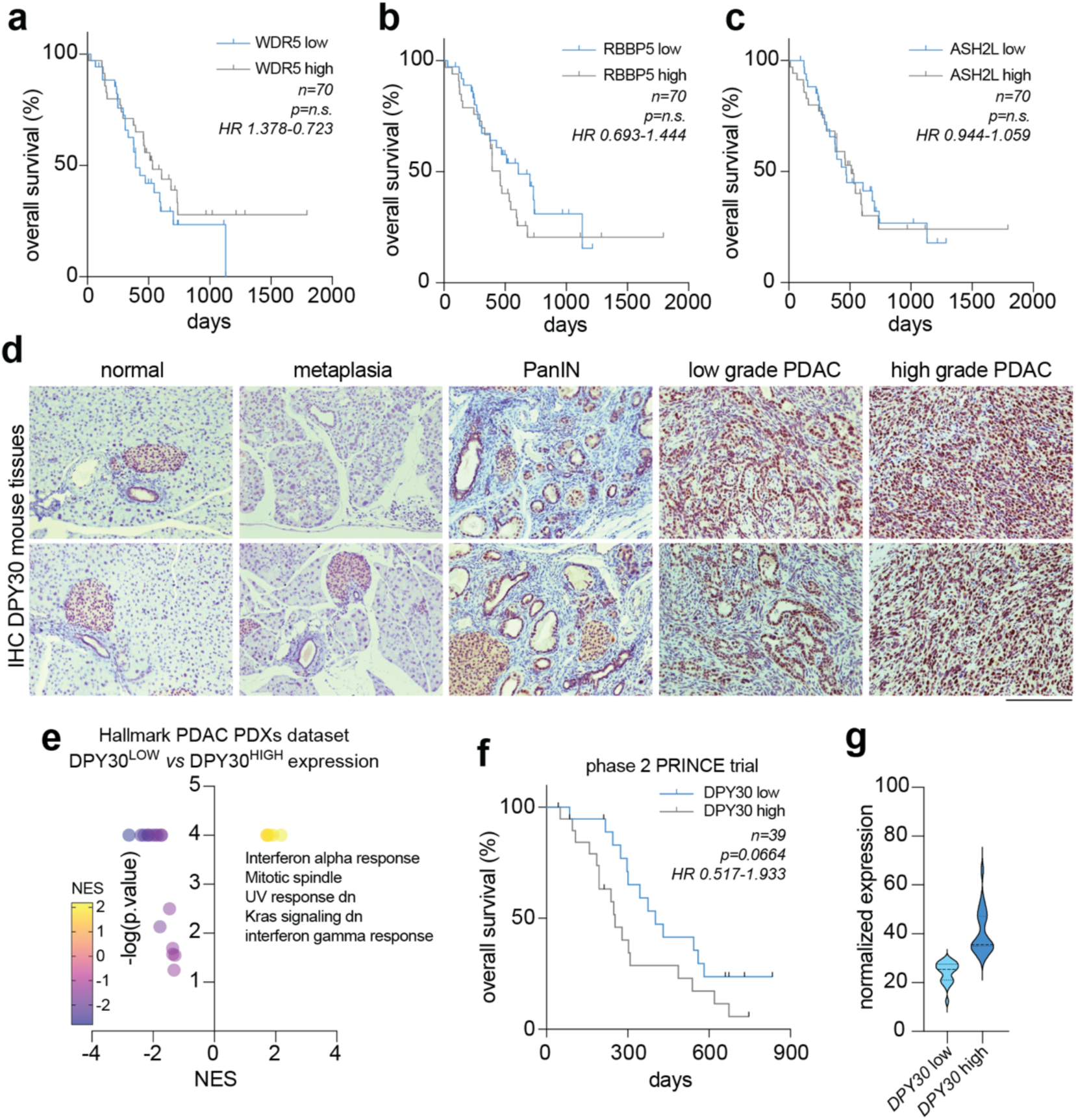
DPY30 correlates with tumor stage and predicts response to ICB treatment. **a-c.** Kaplan−Meier survival curves of TCGA PDAC patients divided according to low- or high-expression of *WDR5* (a), *RBBP5* (b) and *ASH2L* (c). **d.** DPY30 immunohistochemical staining in normal pancreas, pancreatic metaplasia and intraepithelial neoplasia (PanIN) and PDAC samples from KPC mouse model. **e.** Volcano plot representing GSEA in high-purity PDAC PDXs samples and represented as low versus high *DPY30* expression. **f.** Kaplan−Meier overall survival curve of patients in the PRINCE trial cohort divided according to low- or high-expression of *DPY30.* **h.** Violin plot showing the median expression of *DPY30* in the patience included in the analysis in f and Fig. 6h. In (a-c, f), statistical significance and Hazard Ratio (HR) were calculated using the Log-rank test.

